# Oxygen Minimum Zones Drive Changes in Deep-sea Coral Species Distribution, Diversity, and Community Structure on Seamounts in the Eastern Tropical Pacific

**DOI:** 10.64898/2026.04.23.719023

**Authors:** Steven R. Auscavitch, Mary Deere, Morgan Will, Odalisca Breedy, Jorge Cortés, Erik E. Cordes

## Abstract

Oxygen minimum zones (OMZs) are among the most significant abiotic environmental gradients found in the ocean. Yet, fine-scale species distribution patterns of organisms inhabiting OMZs are still spatially limited, hindering our understanding on how these oceanographic features influence species diversity and community structure. Cold-water corals are ecologically important habitat-forming species that are often considered to be sensitive to low seawater dissolved oxygen concentration and thus likely to be useful indicators for exploring change in megafaunal abundance and biodiversity across the OMZ. In the eastern tropical Pacific Ocean, widespread oxygen minimum zones and oxygen limiting zones encompass several thousand square kilometers of area and span several hundred meters of the water column, but typically are strongest between 300-700 m depth. A January 2019 cruise aboard the R/V *Falkor* using the ROV *SuBastian*, conducted video transects along 7 seamounts between the Costa Rica Margin and Isla del Coco, as well as within one submarine incised canyon on the north side of Isla del Coco. In this study, we analyzed survey data for patterns in cold-water coral species distribution, diversity, and coral community structure relative to abiotic oceanographic variables in order to gain biogeographic insights to this area. Across all sites, we identified 3675 coral occurrences and 75 unique morphospecies between 177-1565 m. Rapid species turnover with increasing depth occurred primarily across the upper (300 m) and lower OMZ boundaries (700 m). Coral assemblages within the OMZ depths were observed to contain distinct groups of species compared to those below at deeper bathyal depths. Stylasterid hydrocorals were disproportionately abundant above and within the OMZ, while octocoral and black coral species dominated in the more well-oxygenated waters below. Coral assemblage diversity and abundance was depressed within the OMZ, but bathyal diversity peaked at intermediate water depths between 1200-1500 m. In addition to assessing the impact of OMZs on coral communities, these results provide unique insights to the abundance, diversity, and environmental drivers of deep-water coral community assembly in a data-deficient locality, thus improving biodiversity metrics and informing marine conservation efforts off Costa Rica. These baseline data are particularly salient in the light of projected expansion and shoaling of eastern tropical Pacific oxygen minimum zones as a result of decreasing ocean oxygen concentrations driven by ocean warming and other climate drivers.

## Introduction

Oxygen minimum zones (OMZs) are geographically widespread water column strata where dissolved oxygen concentrations (herein defined as dissolved oxygen concentrations < 22 µmol/L as described by Levin, 2003) decline to hypoxic or anoxic levels resulting in significant effects on deep-sea biota (Levin, Huggett & Wishner, 1991; Levin, 2003; Stramma et al., 2010b; Guilini, Levin & Vanreusel, 2012; Neira et al., 2018). In the eastern tropical Pacific, OMZs are widespread with the geographic distribution and intensity of oxygen concentrations a function of regional oceanographic processes coupled with aerobic respiration by marine organisms in the water column (Karstensen, Stramma & Visbeck, 2008). Specific to the eastern tropical Pacific off Costa Rica, reduced deep-water ventilation in the area of the Costa Rica Dome results in a relatively thick, widespread, and persistent layer of low oxygen between 300-900 m (Karstensen, Stramma & Visbeck, 2008). Environmental heterogeneity resulting from OMZs has been identified as enhancing regional biodiversity in the deep ocean as well as a source of endemic species (Gooday et al., 2010). Persistent OMZs through time are likely to be important boundaries to species distribution, as well as hypothesized barriers to gene flow within populations of deep-ocean organisms (Rogers, 2000; Gooday et al., 2010). Looking into the future, the negative effects of OMZ expansion due to climate change drivers are expected to be strongest within tropical and low-latitude environments (Stramma et al., 2010b; Schmidtko, Stramma & Visbeck, 2017; Sweetman et al., 2017).

Global modelling and experimental efforts have indicated that many benthic megafaunal groups, and particularly cold-water corals (CWC), are highly sensitive to seawater dissolved oxygen concentration (Dodds et al., 2007; Tittensor et al., 2009; Fink et al., 2012; Lunden et al., 2014). For stony corals (Scleractinia) specifically, dissolved oxygen concentrations are predicted to exclude most species from seamount OMZs worldwide (Tittensor et al., 2009). Nevertheless, some examples, like the cosmopolitan reef-building coral *Lophelia pertusa* have been recently found persisting under hypoxic conditions along the Angola margin in the South Atlantic (Hebbeln et al., 2020). Other macrofaunal and megafaunal groups, like benthic foraminifera (Xenophyophores), sponges, and bryozoans (Hanz et al., 2019), are also more likely to be tolerant of dysoxic or hypoxic conditions (Gooday et al., 2010). Examples of benthic fauna tolerating and even thriving in low-oxygen conditions are beginning to become more prevalent, and the mechanisms more clear as additional environmental drivers, like tidal mixing, internal waves, and enhanced surface-derived food supply, are characterized with respect to deep benthic communities (Hanz et al., 2019; Puerta et al., 2020).

### Seamount oxygen minimum zones

Seamounts are large, high-profile (extending > 1000 m above the surrounding abyssal plain) seafloor features that are widely known to contain unique habitat mosaics and diverse biological communities (Rogers, 2018). While the effects of OMZs are more widely-studied on continental margins (Levin et al., 2000, 2002; Helly & Levin, 2004; Jeffreys et al., 2012; Neira et al., 2018; Hanz et al., 2019), the role of OMZs in structuring both pelagic and benthic environments on seamounts are limited to a few case studies (Helly & Levin, 2004). At one relatively well-studied seamount in the eastern North Pacific Ocean off Mexico, Volcano 7 (Wishner et al., 1990, 1995; Saltzman & Wishner, 1997a), the effect of the oxygen minimum zone has resulted in significant zonation effects on both pelagic and benthic communities (Levin, Huggett & Wishner, 1991; Levin, 2002). Macro- and megabenthic organisms were largely absent from the seamount summit (730-750 m) where oxygen concentrations were lowest (0.08-0.09 ml/l), followed by a rapid increase in abundance with increasing depth across trophic levels from fishes to corals and sponges. (Wishner et al., 1990). In the Pacific off central Chile, CWC communities identified by trawl bycatch on one seamount summit were identified as being both dense and diverse, composed of black coral and soft coral species in the comparatively well-oxygenated intermediate waters underlying the OMZ (Cañete & Häussermann, 2012).

In the pelagic environment above and near seamounts, mid-water planktonic species abundances and diversity are also strongly influenced by OMZs (Saltzman & Wishner, 1997a). For example, zooplanktonic species, like the copepod *Rhincalanus nasutus*, have been observed above and below but not within OMZs in the vicinity of Seamount 7 in the eastern Pacific (Saltzman & Wishner, 1997b). The role that OMZs have in structuring deep-water zooplanktonic communities is important both as a driver of mid-water species diversity and abundance (Wishner et al., 2018; Wishner, Seibel & Outram, 2020), but also potentially as a driver of change in food supply for benthic suspension-feeding organisms, including corals (Gage & Tyler, 1992).

### Deep-sea Biodiversity Conservation in Costa Rica

In the eastern Pacific, CWCs are important sources of biodiversity in the deep ocean. Structures formed by coral colonies have been found to provide habitats for invertebrates (Cairns, 1991, 2007, 2018a) and fish fauna alike (Auster et al., 2016). Cocos Island (Spanish: *Isla del Coco*), a national marine park and UNESCO World Heritage Site (Cortés, 2016; Claudino-Sales, 2019), is already known as an important reservoir of shallow-water marine biodiversity (Cortés, 2012). Recent cold-water coral species discoveries have given new motivation to understanding benthic biodiversity in Costa Rican waters (Opresko & Breedy, 2010; Breedy, van Ofwegen & Vargas, 2012; Cortés, 2012; Breedy et al., 2019). Furthermore, mesophotic and shallow-water seamounts, like Las Gemelas, are suspected to host deep-water fish and invertebrate species that remain undescribed and communities that are uncharacterized (Breedy & Cortes, 2008; Starr et al., 2012; Starr et al., 2012).While several protected areas exist off the continental margin of Costa Rica (Alvarado et al., 2012) as well as around Cocos Island (Cortés, 2012; Arias et al., 2016), many of the seamounts off Costa Rica remain external to protected boundaries. Further exploration of these seamounts and characterization of benthic communities and substrate types will aid in better informing conservation strategies for these deep-sea environments.

The presence of well-developed oxygen minimum zones are also important features for the deposition of commercially-valuable metal-rich mineral crusts on seamounts (Halbach & Puteanus, 1984; Halbach et al., 1989). Oxygen minimum zones provide metallic ions to the water column which precipitate onto the seafloor on the order of 1-5 mm per million years (Halbach et al., 1989; Hein et al., 2013). Seamounts and other submarine features along the Cocos Ridge have been highlighted as potential areas containing Co-rich mineral crusts (Miller et al., 2018).

With growing trends towards exploration and exploitation of deep-sea mineral resources (Miller et al., 2018), it is important to understand the abundance and diversity of species inhabiting these depths as to better conserve unexplored or underexplored deep-sea habitats from the potentially destructive influence of mining practices (Niner et al., 2018).

### Objectives and hypotheses

In this study, we aim to characterize the distribution, diversity, and community structure of CWCs across abiotic gradients across the Pacific Costa Rica seamounts extending from the continental shelf to the vicinity of Cocos Island, Costa Rica. The effect of a strong, persistent OMZ in this area is likely to be apparent in the zonation patterns of coral communities across the bathyal depth range. Specifically, we aim to test the hypothesis that CWC assemblages within the OMZ are distinct from those in the waters above and below the OMZAs water mass boundaries are also important drivers of change in coral communities across depth, we will test for differences in CWC community structure among eastern Pacific water masses. As deep-sea communities are expected to be subject to significant changes resulting from global climate drivers (Brito-Morales et al., 2020), establishing baseline datasets for understanding how benthic communities might respond to expansion and intensification (Stramma et al., 2010b) of oceanic OMZs is critical.

## Materials & Methods

### Study Area and ROV Surveys

Surveys were conducted in January 2019 on the R/V *Falkor* using the ROV *SuBastian* o seamounts on the Cocos Plate as well as off Cocos Island proper (Fig. 1). In total, 8 sites were surveyed, 7 seamounts (including Las Gemelas) and 1 site classified as an incised canyon, Cocos Canyon, spanning a total depth range from 179-1614 m (Table 1). A total of 80 hours of seafloor video were assessed among all the locations encompassing 10.5 linear km of seafloor. All surveys were conducted upslope along each feature, perpendicular to the bathymetric contours, typically at an over-ground velocity of 0.25-0.5 kt (0.1-0.25 m/s), stopping only for sampling events or brief detailed imagery. Onboard conductivity, temperature, and depth sensors (SeaBird FastCAT SBE49 CTD) and oxygen optode (Aandera O2 optode 4831) logged continuously in UTC time along the length of each dive profile. Seafloor position was logged using an ultrashort baseline (USBL) navigation sensor (Sonardyne WMT 6G).

**Figure 1:**
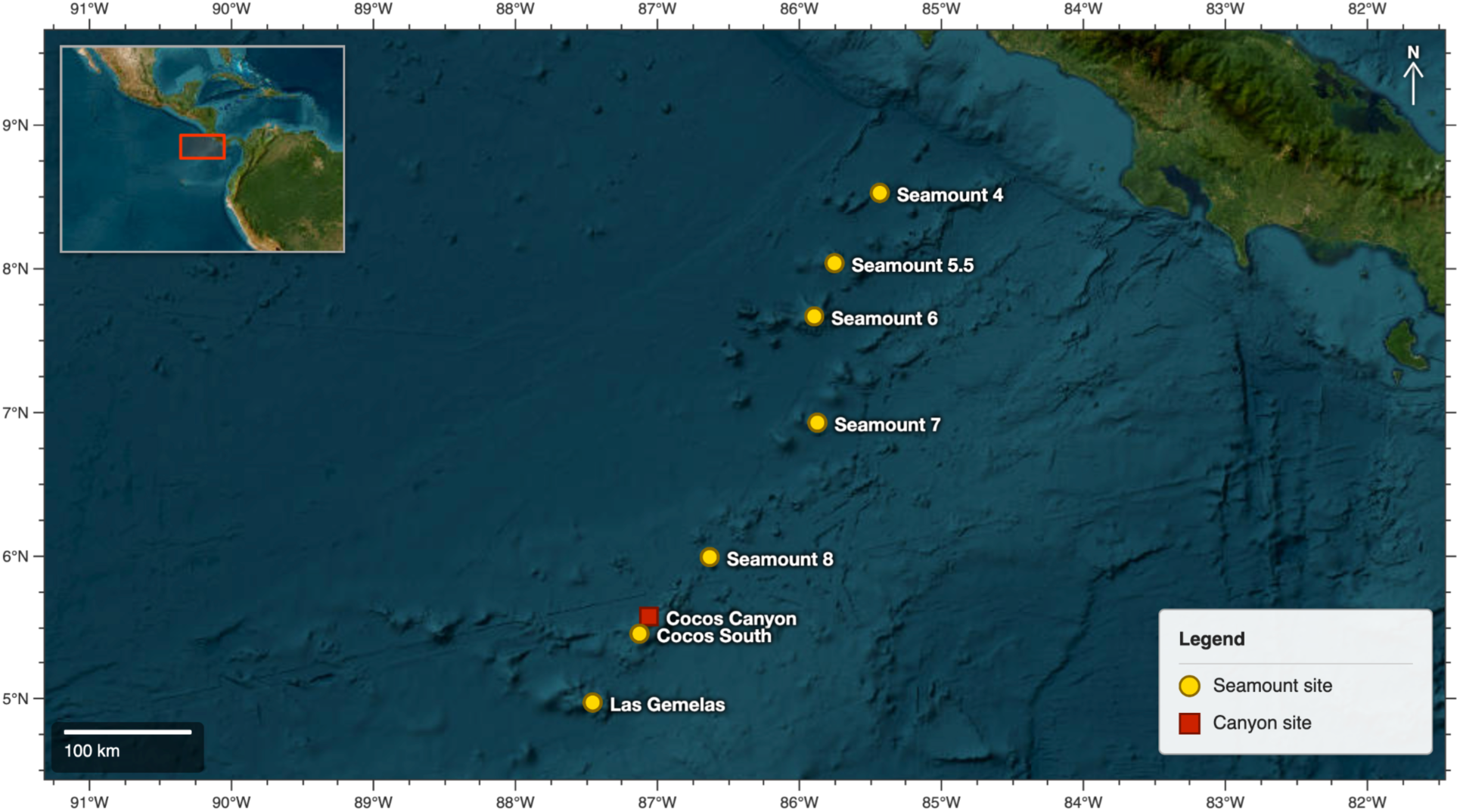
Overview of the study area with locations of seamounts and Cocos Island. Yellow circles represent seamount sites and red squares represent an incised canyon site off Cocos Island.

**Table 1:** Summary statistics for dives conducted during FK190106 along the seamounts and near Cocos Island.

| Dive | Site | Start Latitude (Degrees) | Start Longitude (Degrees) | End Latitude (Degrees) | End Longitude (Degrees) | Bottom Time (hh:mm) | Depth Range (m) | Linear Distance (m) |
| --- | --- | --- | --- | --- | --- | --- | --- | --- |
| S0221 | Seamount 5.5 | 8.04954 | -85.75857 | 8.05072 | -85.77417 | 10:19:00 | 777-1614 | 1721 |
| S0222 | Seamount 7 | 6.93338 | -85.88089 | 6.91501 | -85.88507 | 10:20:00 | 866-1265 | 2092 |
| S0223 | Cocos Canyon | 5.58535 | -87.06810 | 5.58114 | -87.06264 | 10:28:00 | 546-976 | 764 |
| S0224 | Coco South | 5.46551 | -87.13686 | 5.46622 | -87.12633 | 10:44:00 | 179-593 | 1167 |
| S0225 | Las Gemelas | 4.97850 | -87.45997 | 4.98710 | -87.44329 | 10:45:00 | 243-670 | 2079 |
| S0226 | Seamount 8 | 5.99908 | -86.64332 | 6.00781 | -86.64915 | 7:35:00 | 1302-1451 | 1164 |
| S0227 | Seamount 6 | 7.67786 | -85.90642 | 7.68012 | -85.91151 | 9:52:00 | 525-662 | 614 |
| S0228 | Seamount 4 | 8.54279 | -85.43900 | 8.55068 | -85.43904 | 10:09:00 | 888-1260 | 878 |

### Water Sampling and Carbonate Chemistry

A combination of water column (CTD rosette) and sea floor (ROV niskin) seawater samples were aggregated from three cruises: AT3713 (2017) and AT4203 (2018) aboard RV *Atlantis* with HOV *Alvin*, and FK190106 (2019) aboard RV *Falkor* with ROV *SuBastian*. For all samples, pH was measured on deck within 1 hour of collection using an Orion 5-star pH meter, followed by preservation in 500mL high-density polyethylene (HDPE) containers by poisoning with 100 µL saturated mercuric chloride and storage in a cool, dark space (Dickson et al., 2007). Associated physical data (pressure, temperature, salinity, and oxygen) were recorded for each collected sample from vehicle mounted conductivity-temperature-depth (CTD) and oxygen sensors. Total alkalinity was then measured for each sample through acid-titration in triplicate following established methods (Lunden et al., 2013; Georgian et al., 2016) and used in conjunction with pH and associated physical data to calculate aragonite saturation level (Ω_arag_) using CO2SYS (Pierrot et al., 2011). Finally, calculated Ω_arag_ values were used to fit a power regression model to predict Ω_arag_ at each community sample location to allow for direct comparisons.

### Video Analyses

Video was annotated from on bottom to off bottom times for coral occurrences along the length of each dive where the vehicle was in contact with the seafloor (< 2 m altitude above the seafloor, except during vertically sloped segments). Periods of time where the vehicle was not moving forward at a constant velocity or when the seafloor was not clearly visible were not assessed. Coral occurrences were tagged with a UTC timestamp and identified to the finest taxonomic level possible based on imagery and associated specimen collections (Supplementary Table 1). For the purposes of this study, coral taxa refer to deep-water species in the Antipatharia (black corals), Scleractinia (stony corals), Octocorallia (sea fans and other soft corals), and Stylasteridae (lace corals). All corals were typically identifiable to a morphospecies if they were above a roughly 5 cm size threshold gauged by on-screen scaling lasers. Additional taxonomic literature and taxonomist expertise were consulted for specimen collection identification. When a species name could not be readily assigned, corals were given a unique morphospecies identifier based on the corresponding collected sample (e.g. Plexauridae sp. S0227-Q9) or another distinguishing identifier (Supplementary Table 1). Consistent morphospecies identifications were carried across all dives.

### Coral community analyses

In order to evaluate changes in community assembly, all dive transects were binned into segments of 100 m depth samples. Each bin contained only consistent morphospecies observations thereby excluding those observations where a morphospecies could not be identified (e.g. Scleractinia spp.). Each sample reflected both the morphospecies abundance as well as diversity. All samples were standardized by the total abundance and 4^th^ root transformed to down-weight the influence of extremely abundant taxa. Environmental variables for each sample were calculated as the average of depth, temperature, salinity, and dissolved oxygen within the depth bin at each survey location. Variables were normalized to the mean (average = 0, standard deviation = 1) in order to reduce the effect of differences in unit scales among them in the statistical analyses.

Sampling effectiveness was evaluated between sites and across depth by sample-based species accumulation and individual-based rarefaction analyses using the R package Vegan 2.5-6 (Oksanen et al., 2015). Patterns of CWC abundance and diversity across depth were evaluated for each sample using several diversity indices. To evaluate change in species richness we calculated exact species richness (S) and rarefied species richness at 100 individuals, Es(100) (Sanders, 1968; Hurlbert, 1971). Diversity patterns were evaluated using two commonly used indices, Shannon diversity (H’ log_e_ ), a sensitive metric to species diversity, and second, a metric of species equitability, Pielou’s evenness index (J’).

Multivariate community analyses were performed in PRIMER7 with PERMANOVA add-on (Anderson, Gorley & Clarke, 2008). Samples were initially compared in a Bray-Curtis similarity resemblance matrix. One-way analysis of similarity (ANOSIM) tests were used to assess for differences in *a priori* groupings related to the OMZ and overlying water masses. In order to evaluate relationships between *a priori* groupings, both CLUSTER with non-metric multidimensional scaling (nMDS) analyses were employed. Similarity profile (SIMPROF) analysis was used to evaluate the significance level of clusters (Clarke, Somerfield & Gorley, 2008). Bootstrap pseudo-replicates and averages for groups above, below, and within the OMZ were also calculated using nMDS ordination for further investigation of differences among sample means (Clarke & Gorley, 2015). Similarity percentage (SIMPER) analysis was used to identify patterns of dissimilarity among factors as well as indicating the relative importance of species abundances to community assembly. Principal coordinate analysis (PCO) was conducted to further assess relationships among coral assemblages, in addition to identifying coral species that are highly correlated with groups of samples.

For establishing relationships between patterns in environmental variability and community structure, the BEST (BIO-ENV) routine was used to identify the strength of correlations between any single one or groups of environmental variables and coral assemblage variation (Clarke et al., 2008). In order to model the role of environmental variables in structuring CWC communities, a distance-based linear model (distLM, specified variable selection) was constructed using biological and environmental data and distance-based redundancy analysis (dbRDA) (Anderson et al., 2008). An Akaike Information Criterion (AIC) was utilized to evaluate model selections (Akaike, 1998).

## Results

### Identification of the OMZ and local water masses

The depth of the oxygen minimum zone was defined as depths of the water column where the dissolved oxygen concentration was < 22 µmol/L (Levin, 2003). Using this parameter, the core of the OMZ was identified between 300-700m (Fig. 2). The lowest seafloor oxygen values were measured with the *SuBastian* CTD and oxygen optode at both Cocos Island and Las Gemelas around 450-500 m. The core depth of the OMZ was found to be slightly thicker and deeper at ship-based CTD rosette casts closer to the Costa Rica Margin and decreased with distance offshore (Las Gemelas, 300-700 m) (Fig. 2). In some instances, dissolved oxygen values observed on the seafloor *in situ* with the ROV CTD unit (as low as 0 µmol/L, i.e. below detection limits of the sensor) were lower than those observed with the downcast CTD profiles (Fig. 2), which were all conducted in deeper waters off of the seamounts.

**Figure 2:**
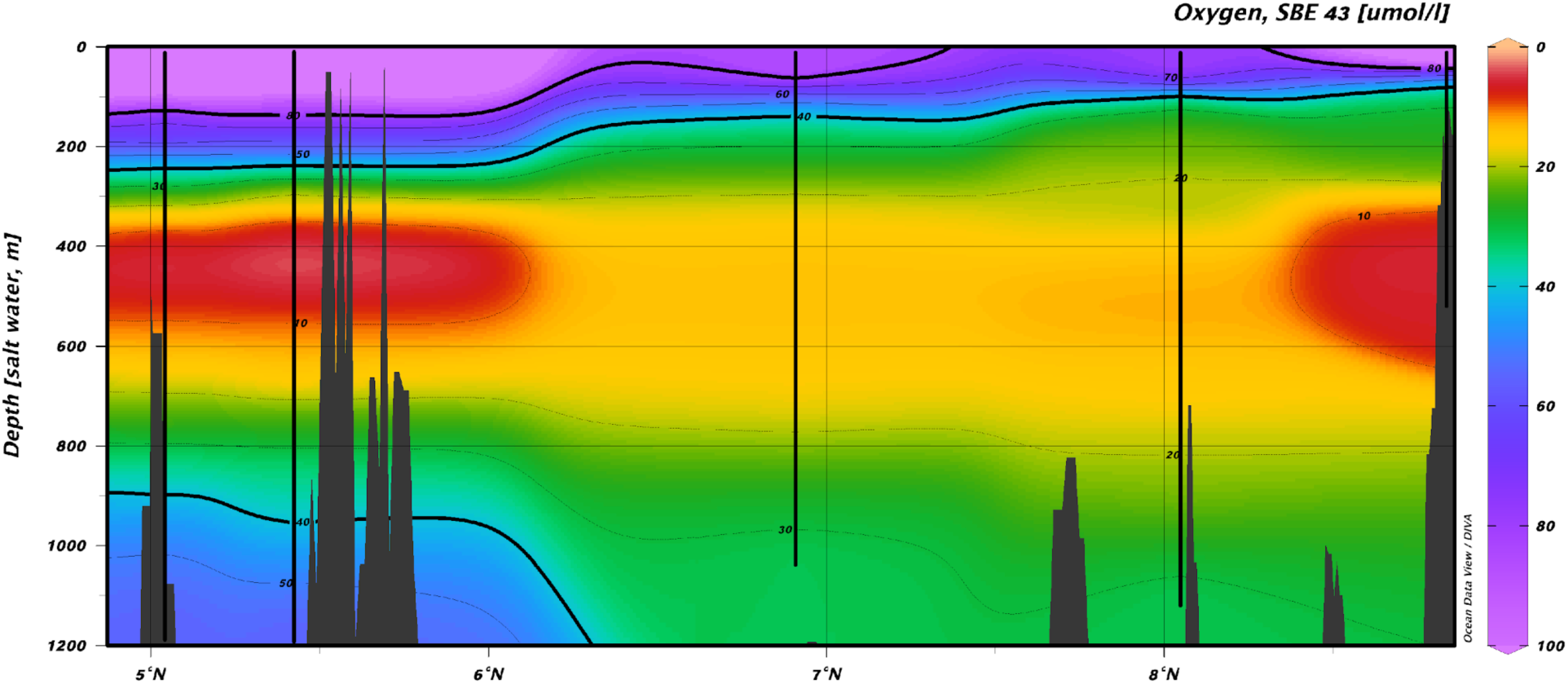
Oxygen profiles along the seamount chain from the continental margin (far right) to Las Gemelas (far left) between 0-1200 m. The color ramp on the right indicates measured values of dissolved oxygen (µmol/l) with cooler colors indicating more oxygen rich waters. Downcast CTD cast locations are represented by bold vertical lines. Bathymetry (GEBCO 2014 2x2 min resolution) is presented as dark grey polygons. Plot was constructed using Data-Interpolating Variational Analysis (DIVA) gridding in Ocean Data View v 5.2 (Schlitzer, 2019).

Water masses were also characterized for the study area of the equatorial eastern Pacific from oceanographic literature that utilized both standard methods (e.g. potential temperature and salinity) as well as geochemical tracers (Fiedler & Talley, 2006; Bostock, Opdyke & Williams, 2010; Peters et al., 2018). Three water masses were identified between 179-1614 m (Table 2).

**Table 2:** Water masses and associated environmental variables in the region of the Cocos Island and nearby seamounts at bathyal depths.

| Water Mass | Depth Range (m) | Temperature (°C) | Salinity (PSU) | Dissolved Oxygen (μmol/L) | Density ($\sigma_t$) | Defining Characteristics | Source Waters | References |
| --- | --- | --- | --- | --- | --- | --- | --- | --- |
| Tropical Surface Water (TSW) | Surface – ~100 | 17-29 | 30.5–34.8 | 90 - 200 | 20–26.2 | Decreasing temperature, increasing salinity. | Surface waters, low salinity resulting from surface freshwater input (e.g. precipitation) | Fiedler & Talley 2006 |
| Equatorial Subsurface Water (ESSW) | ~100–200 | 12-17 | 34.9–34.5 | 50 - 100 | 26.0–27.0 | Sub-thermocline salinity maximum | Small, local water mass of unclear origin, similar to 13°C Water | Fiedler & Talley 2006; Peters et al 2018; Evans et al 2020 |
| Equatorial Pacific Intermediate Water (EqPIW) | 200–1500 | 2-12 | 34.5–34.6 | 0–75, but also up to 100 | 27.3–27.6 | Decreasing salinity, oxygen minimum layer. Sub-pycnocline salinity minimum. | Near Costa Rica, this water mass is a mixture of AAIW (core restricted to 20°S of the equator) and NPIW (from the coast northward to California). | Fiedler & Talley 2006; Bostock et al 2010 |
| North Pacific Deep Water (NPDW) | 1500 – ~2300 | 1–1.8 | >34.6 | >100–125 | 27.6–27.8 | Increasing oxygen, increasing salinity, increasing density. | North Pacific, largest water mass in the North Pacific Ocean. | Fiedler & Talley 2006 |
| Lower Circumpolar Water (LCPW) | >2300 | <0.85–1 | 34.7 | ~200 | >27.8 | Increasing density, increasing O<sub>2</sub>. | Antarctic front | Fiedler & Talley 2006 |

The shallowest, Equatorial Subsurface Water (ESSW) (sometimes referred to as 13°C Water (Evans et al., 2020)), was characterized by a sub-thermocline salinity maximum and rapidly decreasing oxygen concentration between 100 to ∼200 m. Underlying this was the largest water mass, by thickness, Equatorial Pacific Intermediate Water (EqPIW), between the depths of 200-1500 m which also contained the OMZ. Equatorial Pacific Intermediate Water in the eastern tropical Pacific can be differentiated into North and South EqPIW by dissolved oxygen concentration where North EqPIW is oxygen poor compared to South EqPIW (Bostock, Opdyke & Williams, 2010). The deepest, North Pacific Deep Water (NPDW), occurred between 1500-2300 m and was defined by increasing dissolved oxygen concentration and increasing salinity with depth. It was also indicated that Lower Circumpolar Water (LCPW) underlies NPDW (Fiedler & Talley, 2006), but insufficient CTD profiles from depths >2300 m were conducted for the 8 sites focused on in this study (Fig. 2).

### Water Column Carbonate Chemistry Analysis

Water samples for carbonate chemistry analyses were collected at depth from 5 to 1886 m (Supplementary Dataset D1). Total alkalinity ranged from 1982.4 to 2444.5 μmol·kg−1, and pH ranged from 7.389 to 8.174 , with minimum pH observed at 1040 m depth. Calculated Ω_arag_ values from Niskin bottle samples ranged from 0.593 at 1040 m depth to 1.196 at 242 m depth (Fig. 3, A). The aragonite saturation state of the water column was modelled using a power function regression according to the following equation: Ω_arag_ = 6.4829 × depth(m)^−0.3235^ (*R*² = 0.873, *p* < 0.001, *n* = 267) (Fig. 3, B). The saturation horizon where Ω_arag_= 1.0 was identified at approximately 323 m, roughly coincident with the upper boundary of the oxygen minimum zone.

**Figure 3:**
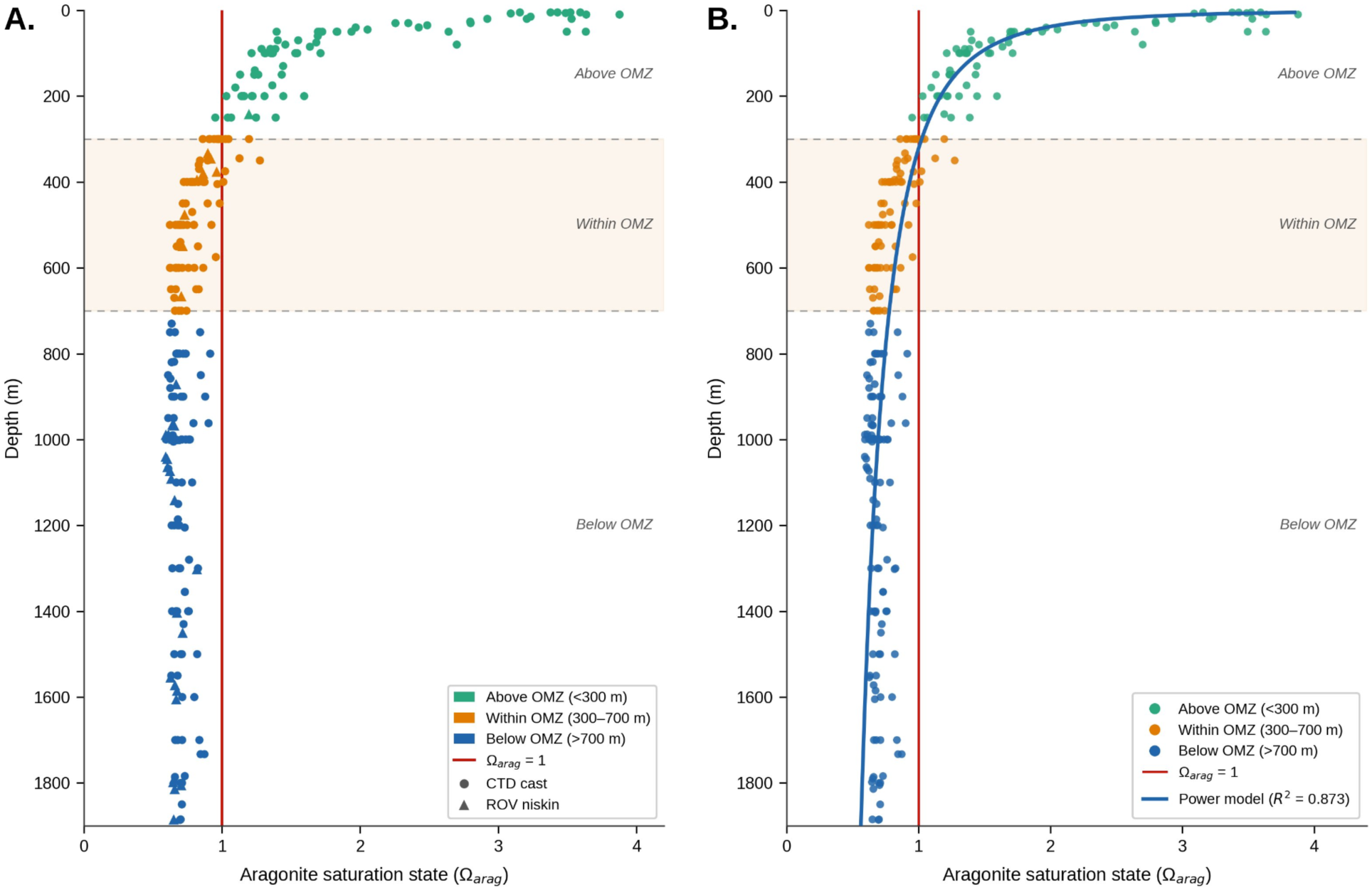
Scatterplots of aragonite saturation states from water samples collected in the area of the Costa Rica continental shelf to Cocos Island from the surface to 1900 m depth (A.). The aragonite saturation horizon (Ω_arag_=1) is indicated in red and the oxygen minimum zone is shaded. Collection method, CTD rosette (closed circles) or ROV Niskin (closed triangle) are indicated. B. Regression of best fit through the saturation state data resulted in the power model Ω_arag_ = 6.4829 × (depth, m)^−0.3235^ (*R*² = 0.873, *p* < 0.001, n = 267).

### Video survey summary

In all, 3675 coral occurrences were recorded between 177 – 1565 m depth across all 8 sites. Of these occurrences, 3386 observations were assigned to 75 identifiable coral morphospecies. (Fig. 4). The remaining 289 observations were assigned to coarser taxonomic groupings based on the observable traits in video imagery (Supplementary Table 1). Of the 75 morphospecies identified, 23% (17 species) were identified as a currently described species based on morphological examination of the associated collection(s) and taxonomic expert consultation (Supplementary Table 1).

**Figure 4:**
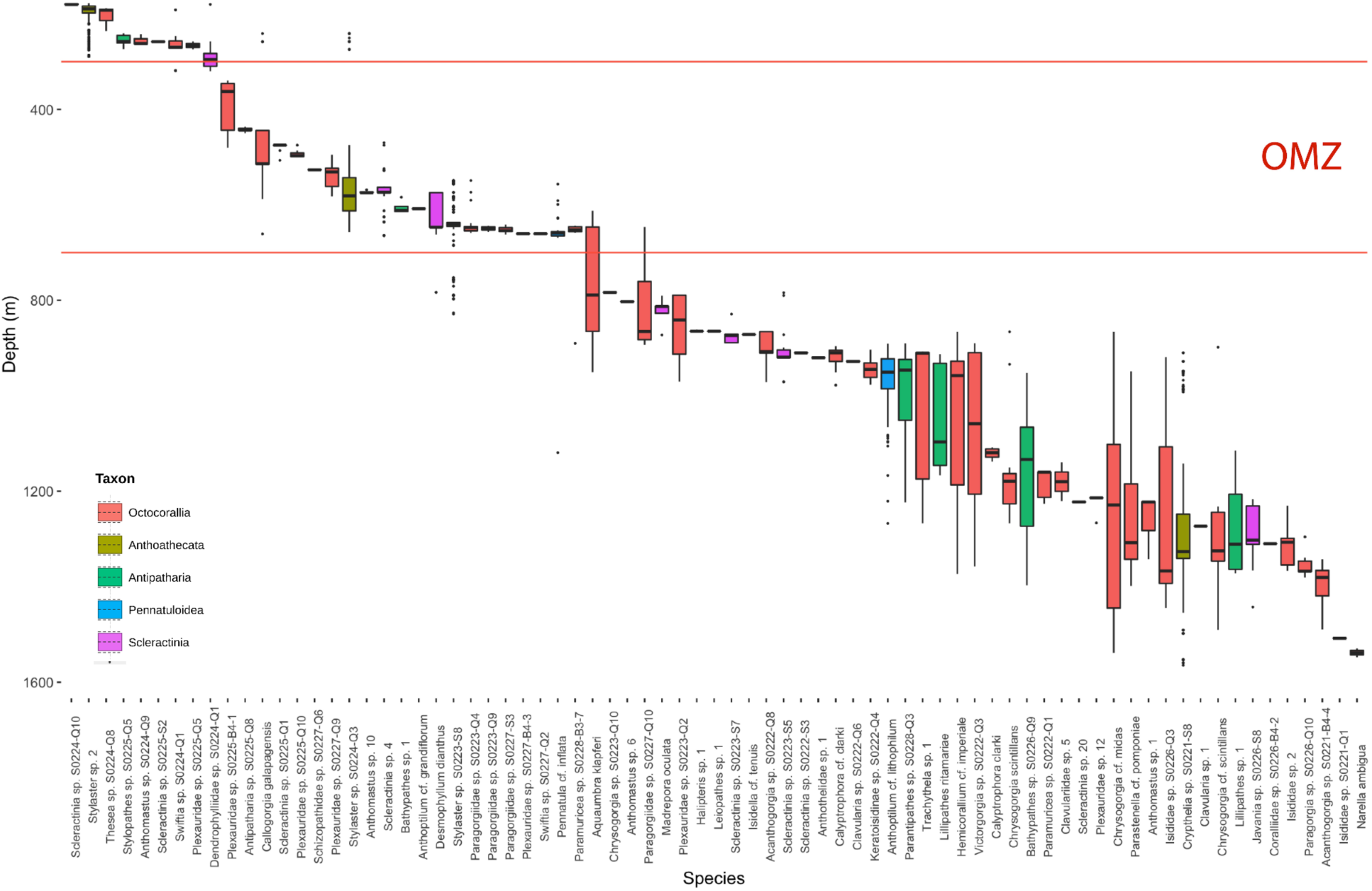
Boxplot of species distribution across depth. Species are arranged left to right by increasing median depth of distribution. Boxes are colored by taxonomic order for each species. OMZ depths (300-700 m) are indicated by solid red lines and bold red text.

Across all depths, the Octocorallia dominated CWC assemblages by abundance (39%) as well as species richness (46 spp.). Inclusive of the Pennatuloidea, the Octocorallia (50 species in total) comprised 55% of all corals observed. Stylasterid hydrocorals followed with 1185 records across 4 species, all within the genera *Stylaster* or *Crypthelia*. In nearly all cases, representatives of each order were observed across the full range of depths explored on the seamounts and Cocos Island (177-1565 m) (Fig. 4). Scleractinians (294 records, 12 spp.) and Antipatharians (101 records, 9 spp.) were present from 179-1508 m but these groups were not as abundant as the Octocorallia and Stylasteridae (Supplementary Table 1). Among the Scleractinia, solitary species (cup corals) were more abundant and occurred across a greater depth range than colonial species (Supplementary Table 1).

### Species Accumulation and Rarefaction Analyses

Species accumulation curves suggest incomplete species inventories at nearly all sites, with the exception of Seamount 5.5 (Fig. 5A). This was the site with the largest depth range surveyed, and species richness was found to approach asymptotic values at approximately 12 species, suggesting a relatively well-sampled community (Fig. 5A). The remaining sites continued to accumulate species at similar rates. Seamount 7 was found to have more rapid species accumulation at lower levels of sampling effort, indicating high species richness at that particular site (Fig. 5A). Individual-based rarefaction analyses indicated the highest species richness at depths below the oxygen minimum zone (Fig. 5B). The transects within the OMZ layer were found to have roughly half the number of coral species as compared to the group of transects immediately underlying (> 700 m). The shallowest layer immediately above the OMZ was the narrowest layer by depth (100-300 m) and coincided with the fewest number of coral morphospecies observed in those transects (Fig. 5B).

**Figure 5:**
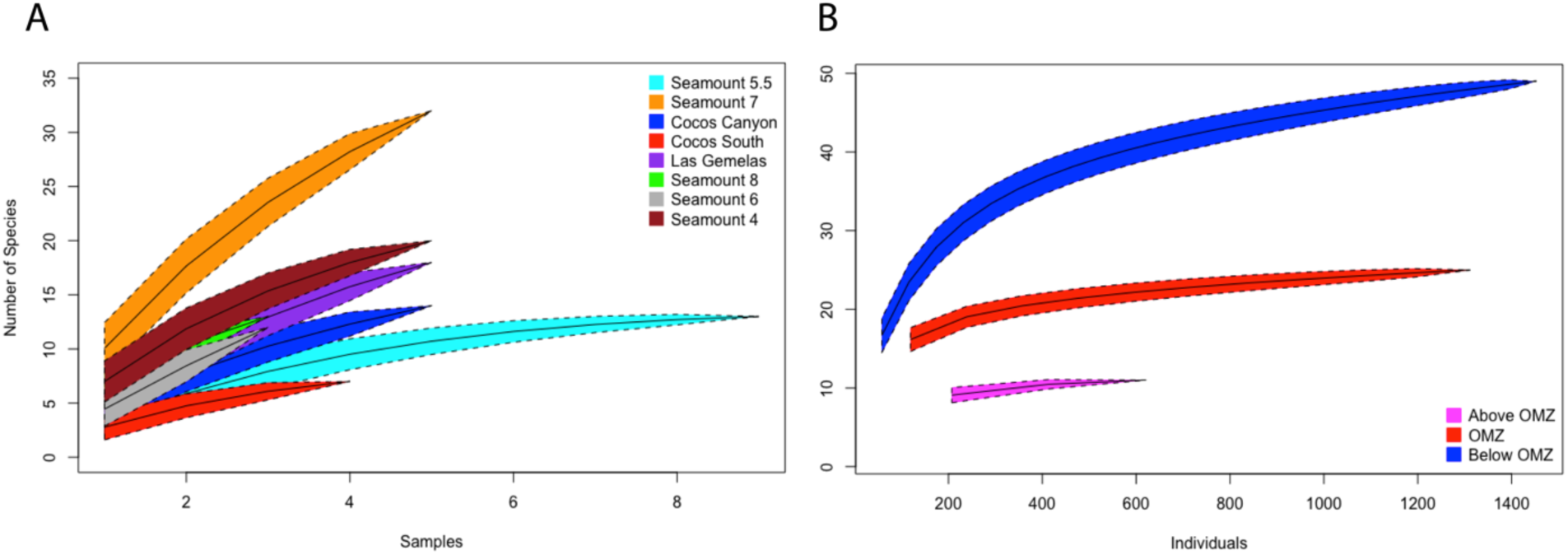
Species accumulation and rarefaction analyses. Permuted (1000 permutations) species accumulation (Coleman et al., 1982) curves are assessed between sites (A). Sample-based rarefaction curves (B) are indicated for each layer of the water column corresponding to depths above (100-300 m), below (700-1600 m), and within (300-700 m) the oxygen minimum zone. For both plots, confidence intervals (95%) are indicated by polygons around the curve bounded by dashed lines.

### Patterns in Coral Abundance and Diversity

Across the entire depth range surveyed, the highest coral abundances were observed within the shallowest depth bin, 100-200 m (Fig. 6A). Within the oxygen minimum zone, mean abundances (the average across seamounts of the abundances in each transect within each depth bin) were lowest at the upper boundary (300-400 m) and peaked along the lower boundary (600-700 m) (Fig. 6A). Mean abundance was generally low (< 30 indiv.) at lower bathyal depths except within the 900-1000 m and 1300-1400 m ranges which were slightly higher. A localized regression model of coral abundances indicated two distinct peaks in abundance, one at the shallowest depth bin and one at the lower boundary of the OMZ (Fig. 6B).

**Figure 6:**
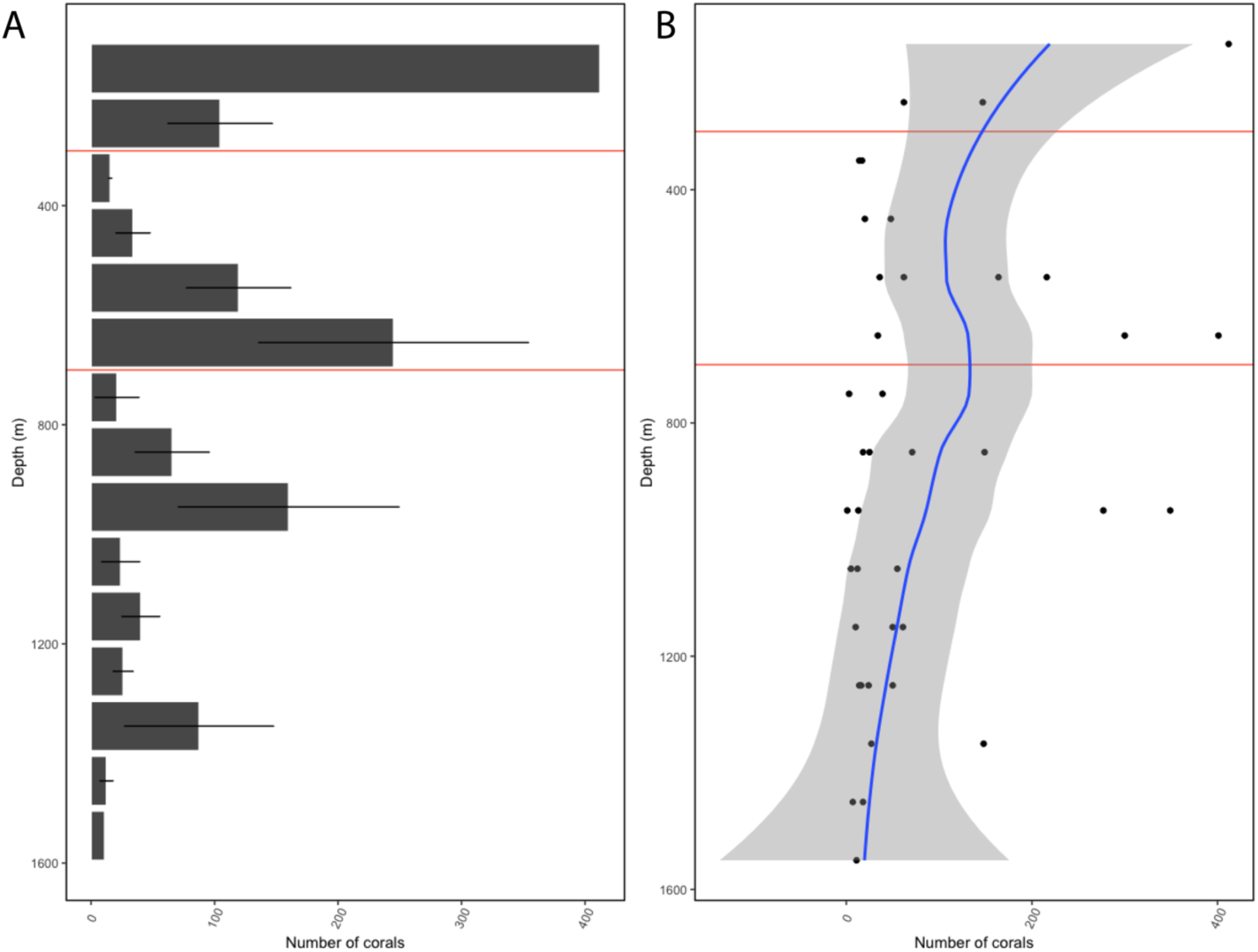
Patterns of coral abundance across depth along the seamount chain. The depth of the oxygen minimum zone is plotted as solid red horizontal lines (300-700 m). (A) Mean coral abundance (+/- 1 Standard Error) is indicated for each 100 m depth bin from 100-1600 m. (B) Scatterplot showing the number of coral records in each sample with a fitted localized regression (loess) model represented by a solid blue line. The shaded region represents the 95% confidence interval for the regression model.

Mean species richness (S) was observed to initially decline with increasing depth reaching an initial minimum in the upper oxygen minimum zone, but then increasing to a maximum around 1200 m (Fig. 7A). The patterns in coral species richness were also reflected in both Shannon diversity (H’) and ES(100) rarefaction indices (Fig. 7B, C). In contrast, mean species evenness (J’) was lowest at shallower depths above the OMZ and increased with depth to maxima just below the OMZ lower boundary (700-900 m) and between 1200-1300 m (Fig. 7D).

**Figure 7:**
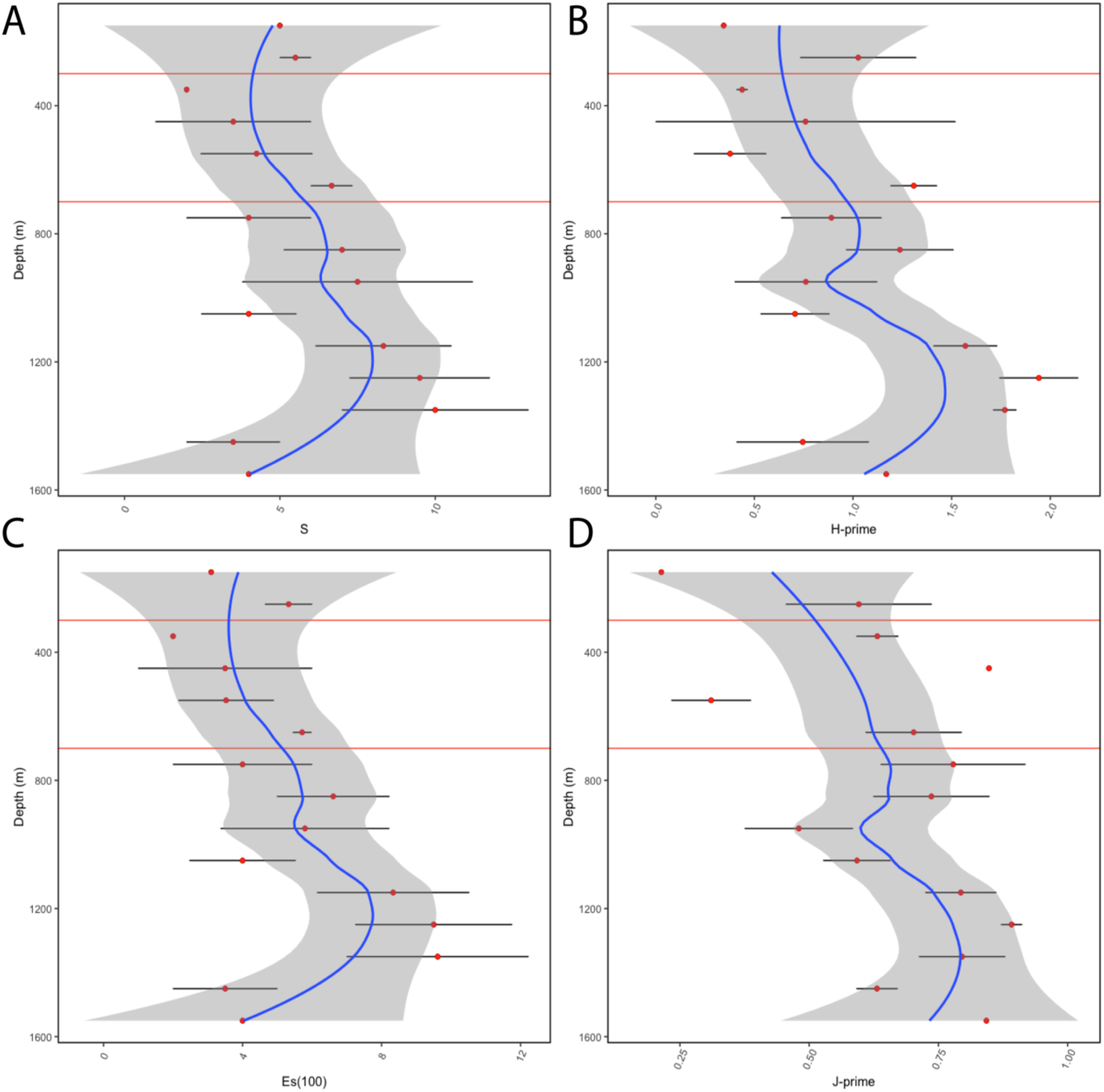
Patterns of coral abundance across depth along the seamount chain. Four diversity indices are indicated: (A) Species richness, S, (B) Shannon Diversity Index (H’ log_e_), (C) Es(100) species rarefaction, and (D) Pielou’s Evenness Index (J’). Points represent mean index values (+/- 1 Standard Error) In each panel the depth of the OMZ is indicated by solid red horizontal lines (300-700 m). For each index, a fitted local regression (Loess) model is represented by a solid blue line. The shaded region represents the 95% confidence interval for the regression model.

### Patterns in Coral Community Structure

An analysis of similarity test (Global R=0.578, p=0.001) between depth strata above, below, and within the OMZ indicated significant differences between assemblages within the OMZ and below the OMZ (R-statistic=0.587, p=0.001) as well as between those above and below the OMZ (R-statistic=0.683, p=0.002) (Supplementary Table 2). The same test did not detect a significant difference between the community above and within the OMZ (R=0.144, p=0.132). A one-way ANOSIM test between assemblages factored by water mass (EqSSW, EqPIW, and NPDW) failed to identify significant differences between any pairwise comparison (Global R=0.152, p=0.16) (Supplementary Table 3).

Permuted SIMPROF tests (999 permutations) of the coral assemblage CLUSTER analysis (Fig. 8) identified 6 distinct groupings of 100 m depth bins at or below the 5% significance level (Fig. 9A). The CWC assemblages showed high similarity above and within OMZ depths (Fig. 8).

**Figure 8:**
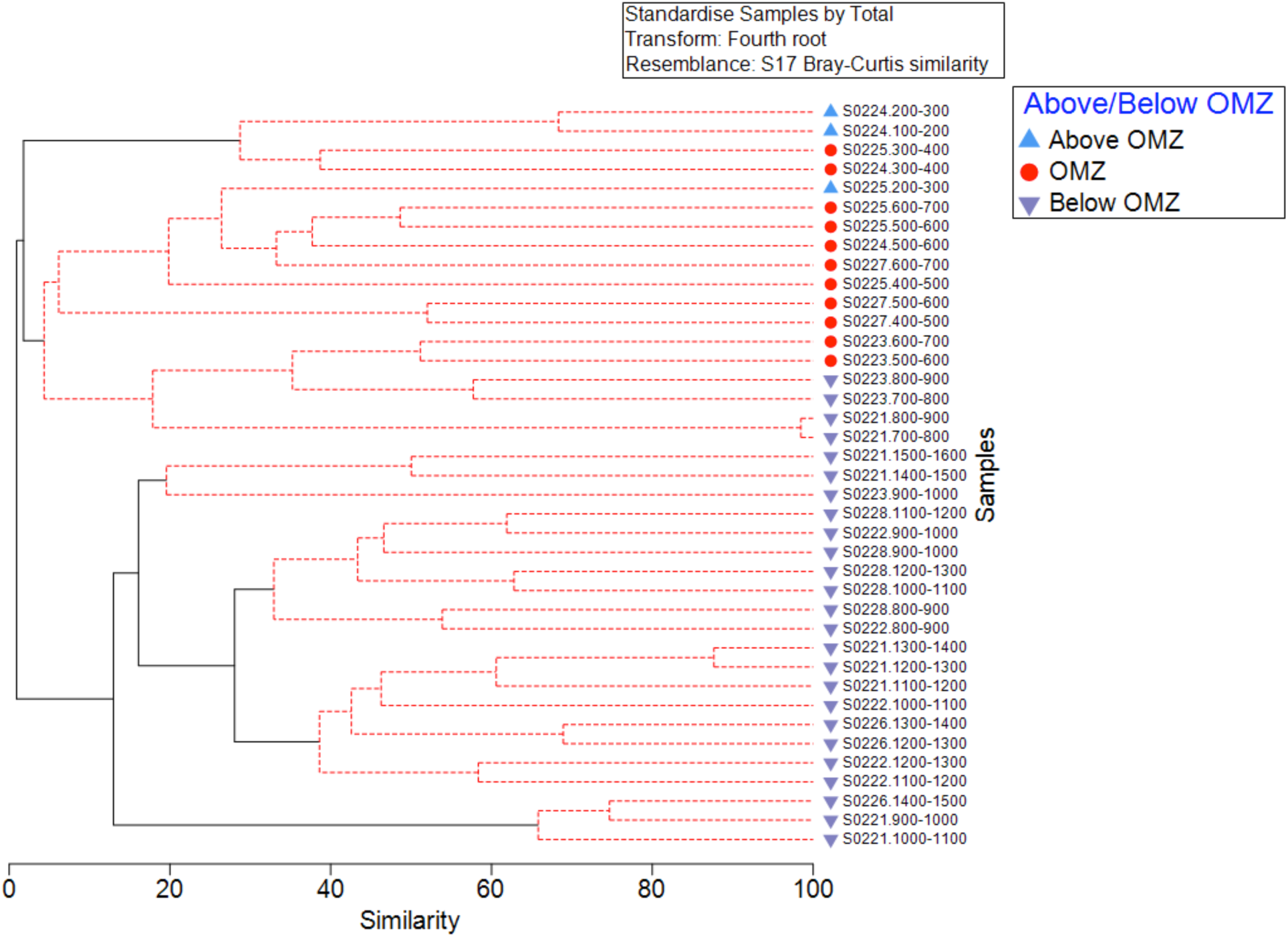
CLUSTER analysis performed on 39 coral assemblage samples from seamount sites. Samples are identified by the dive number and depth range (e.g. S0226.1200-1300) and symbols indicate within, above, or below the depth of the OMZ. Similarity profile analysis (SIMPROF) is indicated by solid black lines for statistically significant clusters (p < 0.05) and dashed red for insignificant clusters.

**Figure 9:**
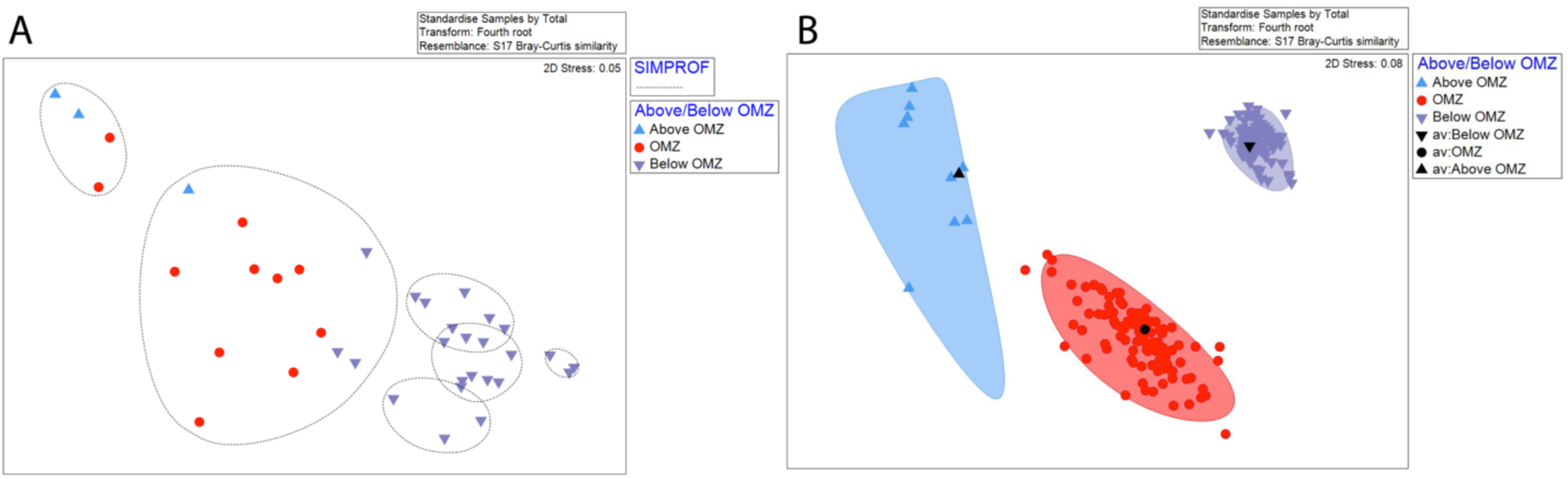
Non-metric multidimensional scaling analyses of 100 m depth sampling units. (A) Each point represents the sum of all transects within a depth bin. These are further indicated as being above, below, and within OMZ. Circles represent statistically significant groupings according to the SIMPROF analysis results indicated in Figure 8. (B) Bootstrap averages analysis of all transects resampled 100 times. Ellipses represent bootstrap regions where 95% of the bootstrap averages fall for each group.

Deeper assemblages occurring below the OMZ were largely clustered into four distinct significant groupings (Fig. 9A). In the bootstrap resampling of samples analysis showed three distinct clusters of transects organized by OMZ factors, with the highest beta diversity in those above the OMZ (Fig. 9B).

Principal coordinates ordination revealed about 33% of the total coral assemblage variation explained by the first two PCO axes (Fig. 10). Two distinct clouds of samples were evident, one grouping above and within the OMZ along PCO axis 2, and a second grouping for samples primarily occurring below the OMZ along PCO axis 1. Vector analysis identified 14 species that were correlated (cutoff R > 0.40) with samples and were driving these patterns in assemblage similarity (Fig. 10).

**Figure 10:**
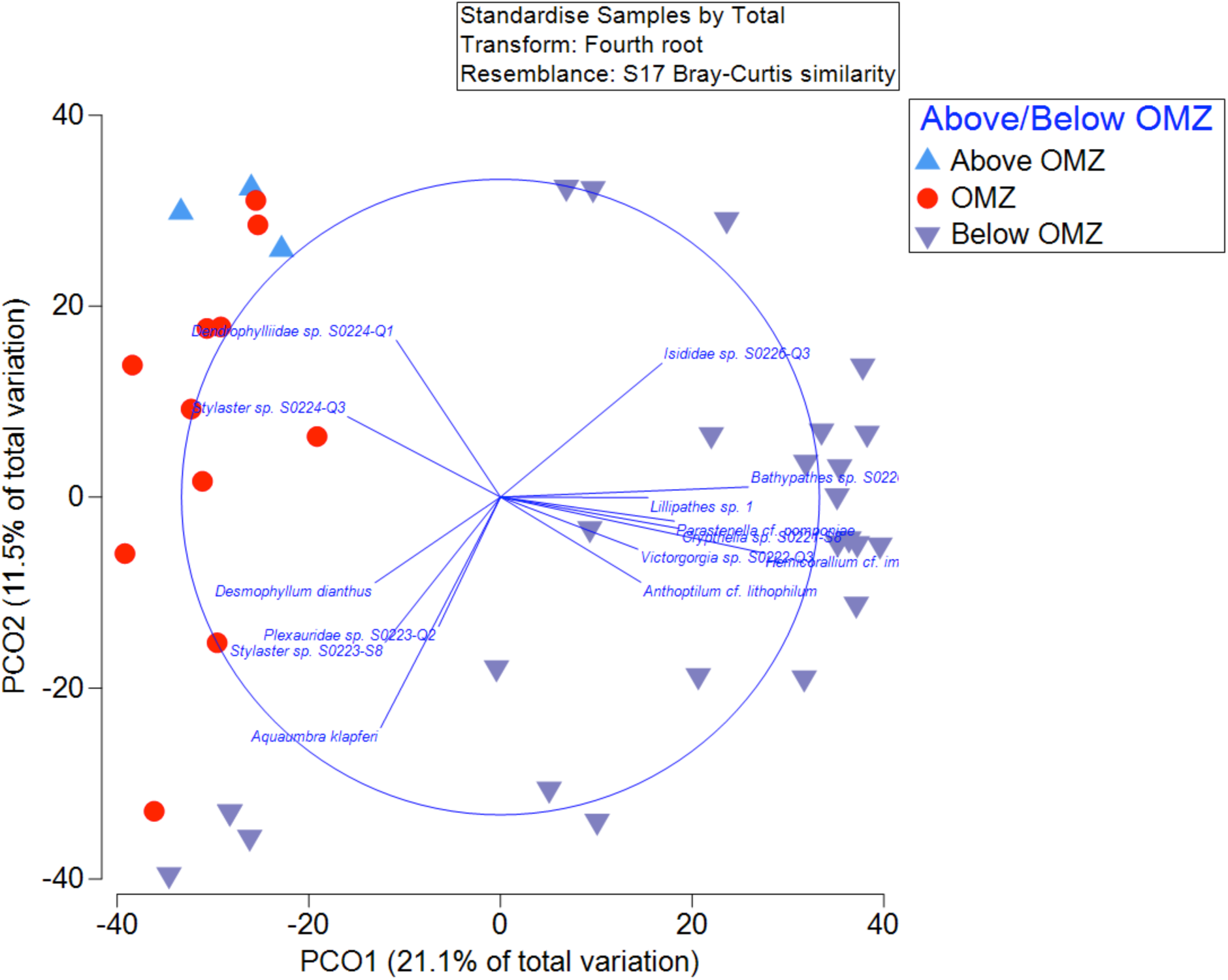
Principal coordinates analysis of 39 samples designated as above, below or within the OMZ. A circle with vector lines indicates any correlations (R= >0.40 cutoff) of species with samples in the PCO space.

SIMPER analysis revealed within-group assemblage similarity was highest at depths above the OMZ (31.7%) (Supplementary Table 4). This was followed by assemblages occurring below the OMZ where the average similarity was found to be 19.7%. Within the OMZ, the average similarity between samples was the lowest at 12% of all three depth strata. As these group similarity values are also indicators of beta diversity, or change in species composition across a gradient, the greatest change in CWC assemblage occurred across the OMZ (300-700 m). Stylasterid corals were identified as having the highest average abundance and being the most representative taxon of the upper two depth strata, above the OMZ and within the OMZ (Fig. 11). Between-group dissimilarity was high (>90-100%) for all pairwise comparisons.

**Figure 11:**
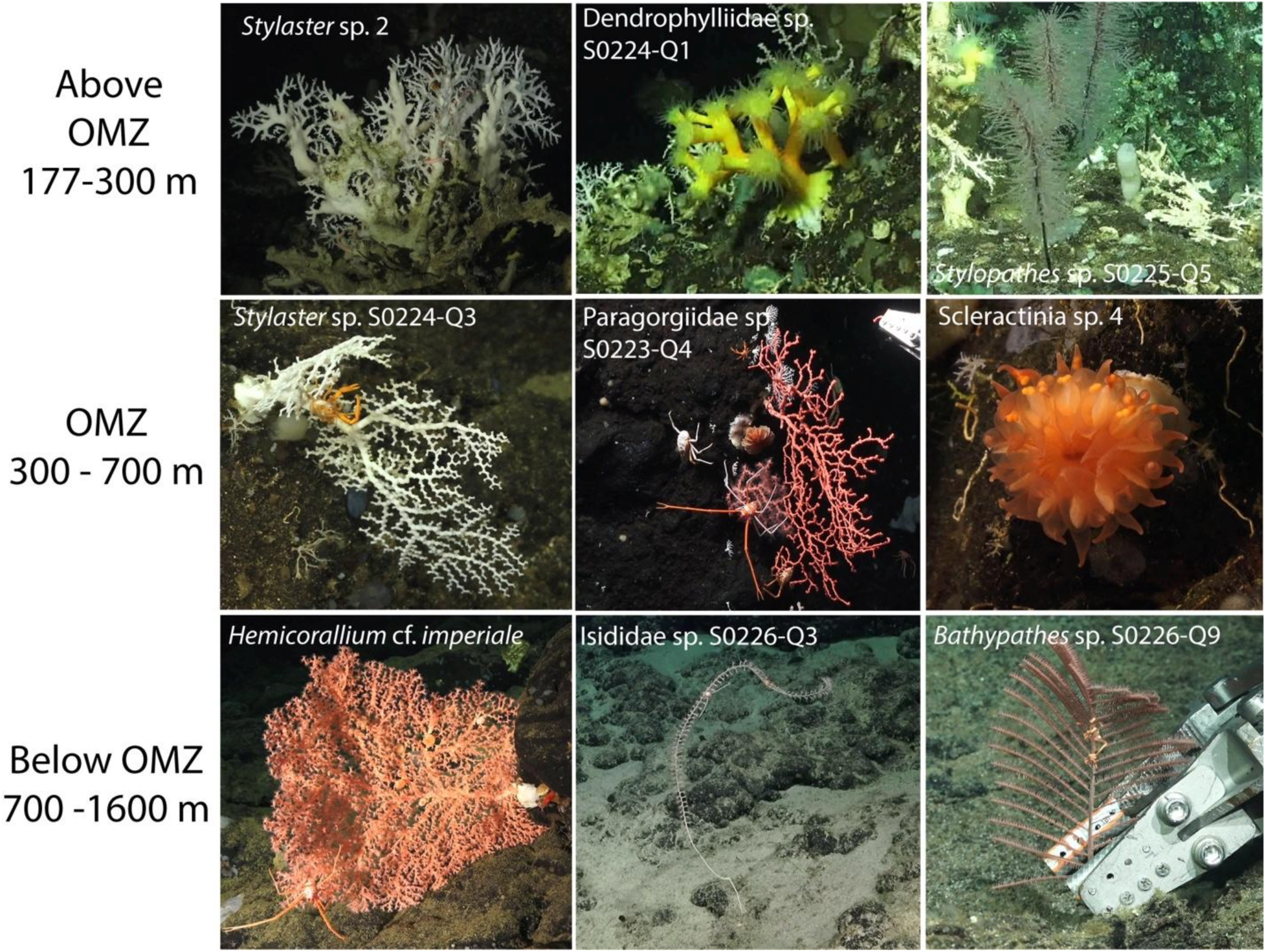
Characteristic species observed within, above, and below the depth of the OMZ along the seamount chain and Cocos Island. The top three species by relative abundance were determined by SIMPER analysis and are arranged left to right in order of decreasing contribution to the total cumulative abundance.

The same analysis also indicated that the shallowest depth layer was characterized by assemblages dominated by stylasterid (*Stylaster* sp. 2), scleractinian (Dendrophylliidae sp. S0224-Q1), and black corals (*Stylopathes* sp. S0225-Q5). The OMZ was composed of a mixed assemblage dominated by a second *Stylaster* sp. (S0224-Q3) as well as soft coral fans (Paragorgiidae sp. S0223-Q4, Plexauridae sp. S0227-Q9) and one solitary coral species, Scleractinia sp. 4 (Fig. 11). At depths below the OMZ, octocorals (*Hemicorallium* cf. *imperiale* Isididae sp. S0226-Q3, and *Aquaumbra klapferi*), followed by black corals (*Bathypathes* sp. S0226-Q9) and one stylasterid, *Crypthelia* sp. S0221-S8, composed at least 70% of the cumulative abundance (Fig. 11).

### Drivers of Change Community Structure across Environmental Gradients

Transitional zones in deep-water coral assemblage composition across depth also closely tracked the aragonite saturation gradient. Stylasterid hydrocorals (Order Anthoathecata), which dominated assemblages above and within the OMZ (0–700 m) (Fig. 4), occurred across a range of Ω_arag_ values from near-saturation to moderately undersaturated conditions (mean Ω_arag_ within OMZ = 0.81 ± 0.14). In contrast, scleractinian corals, which contributed most strongly to assemblages above the OMZ (mean Ω_arag_ = 1.94 ± 0.86), declined sharply within the OMZ and below the saturation horizon (∼323 m). Octocorals and Antipatharians, which dominated assemblages below 700 m, occurred almost exclusively under persistently undersaturated conditions (mean Ω_arag_ = 0.69 ± 0.07), reflecting a hypothetical tolerance of carbonate undersaturation across deep Pacific octocoral communities.

The BEST (BIO-ENV, Spearman rank correlation based on a Euclidean distance resemblance matrix) analysis conducted with depth, temperature, salinity, and dissolved oxygen indicated that temperature alone had the highest correlation (Rho=0.692) among any combination of environmental variables. Additional inclusion of variables in any combination resulted in slightly lower predictive capability (e.g. depth, temperature, Rho = 0.670).

Patterns in coral community structure as explained by oceanographic environmental variables were examined in a distance-based linear model and distance-based redundancy analysis using depth, temperature, salinity, aragonite saturation state (Ω_arag_), and dissolved oxygen variables (Fig. 12). The DistLM marginal tests reported explanatory significance in each of the variables. However, in sequential tests, Ω_arag_ (AIC=319.13, p=0.363) failed to find significance in contrast to depth, temperature, oxygen, and salinity (Supplementary Table 5). In the marginal tests, depth was identified as the variable of best fit explaining 16% of the assemblage variation alone (sequential test, AIC=321.41, p=0.001). The inclusion of Ω_arag_ in a five-variable specified model did not improve model fit relative to a four-variable solution where Ω_arag_ was excluded (AIC = 319.13 versus 318.39 for the four-variable model; R^2^ = 0.356).

**Figure 12:**
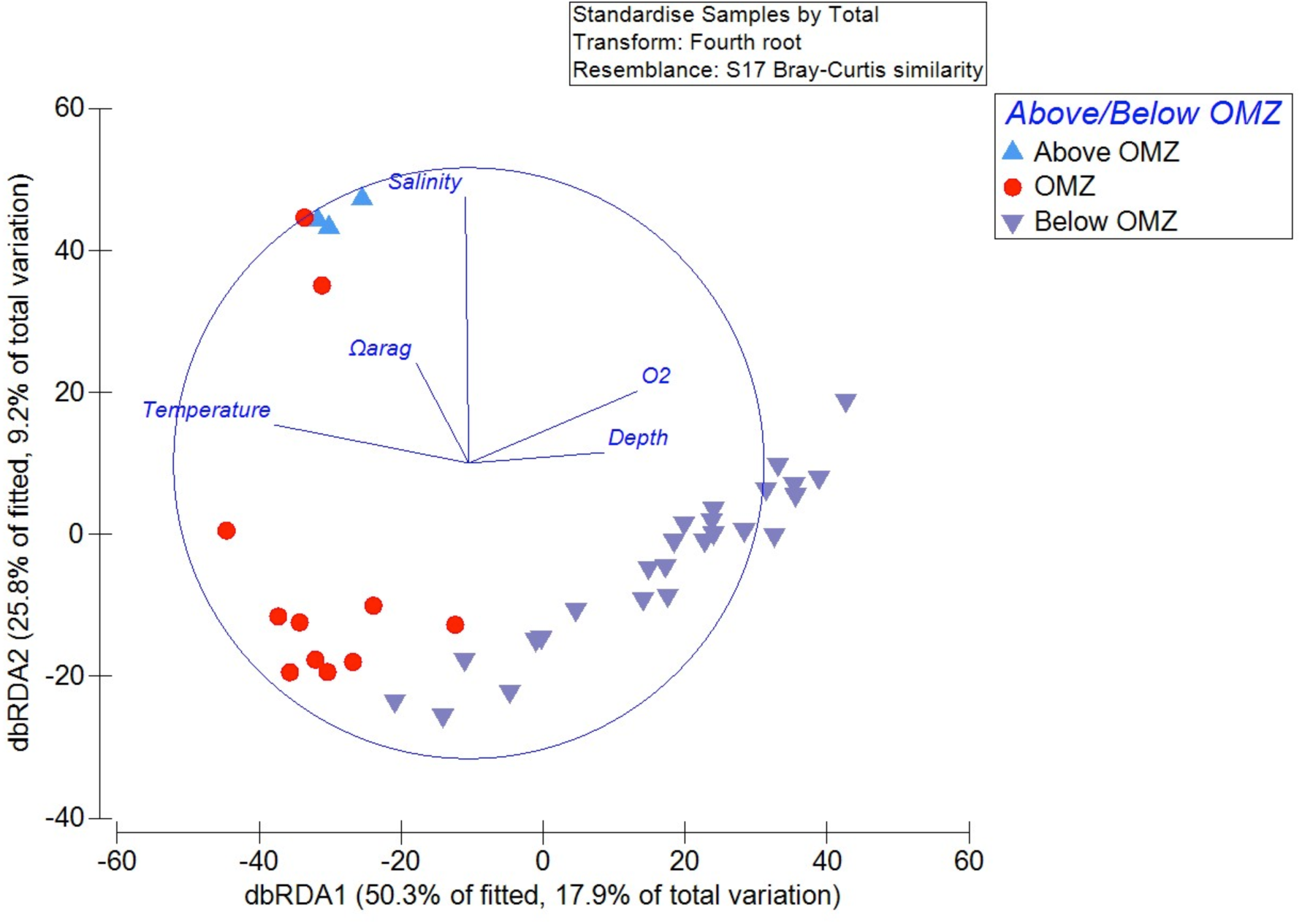
Distance-based linear model (distLM) and redundancy analysis (dbRDA) of coral assemblages and oceanographic variables. Points represent one 100 m depth assemblage. Axes titles are the results of a distance-based redundancy analysis with percent variation explained by the best fitted model and total variation. The dbRDA circle vectors identify the degree of correlation of each environmental variable with samples in the distLM space. The length of each vector is indicative of the strength of the correlation with the circle perimeter denoting a correlation of R=1).

Redundancy analyses indicated that the first two dbRDA axes combined explained 76.1% of the model-fitted variation and 27.1% of the total variation in coral communities (Fig. 12).

Multiple partial correlations revealed that dbRDA axis 1 was most highly correlated with temperature (r=0.66), dissolved oxygen (r=-0.57), and depth (r=-0.46), reflecting the OMZ oxygen and temperature gradient. In contrast, dbRDA axis 2 was strongly and negatively correlated with salinity (r=-0.899) (Supplementary Table 5). Salinity and temperature had the strongest vector associations with the shallowest assemblages (i.e. within and above the OMZ), while Ω_arag_ had a secondary vector association (r=-0.34) along dbRDA axis 2, oriented in the same direction as salinity (i.e. toward the shallowest assemblages) (Fig 12). The vectors for dissolved oxygen and depth were most strongly associated with assemblages below the depth of the OMZ (Fig. 12)

## Discussion

In this study we sought to characterize the distribution, diversity, and community structure of deep-water corals across environmental gradients off Pacific Costa Rica. More broadly, the results of this work provide critical baseline knowledge of deep-water coral species and the communities they compose in the eastern tropical Pacific. In all, 3386 observations of corals were identified among 75 morphospecies between 177 – 1565 m, adding important volume to local and regional species inventories in the eastern tropical Pacific. As some smaller taxa like stylasterids and solitary cup corals often have few distinguishing characteristics to always place them into consistent morphospecies, it should be noted that these abundances represent a minimum value and the likelihood of identifying more individuals and more species increases with additional survey and collection effort.

Octocorals dominated the coral fauna by abundance and species richness at bathyal depths with 55% of all coral observations and 67% of all identified morphospecies. However, stylasterids were disproportionately abundant within the depths of the OMZ and shallower. The elevated abundance of stylasterids in the OMZ may be related to adaptations to low oxygen, but it may also be due to a degree of physiological tolerance to carbonate undersaturation facilitated by their putative high-Mg calcite skeletal mineralogy (Cairns & MacIntyre, 1992). Scleractinians showed their lowest abundance within the OMZ, but then increased again at the deeper depths including *Madrepora oculata*, a structure forming colonial species. If their distribution was simply controlled by Ω_arag_ then their abundance should have stayed low consistent with dissolution pressures exerted on aragonitic corals in undersaturated waters. Their increased abundance below the OMZ suggests that it is oxygen rather than Ω_arag_ that primarily controls their distribution in the Eastern Tropical Pacific.

In general, there were greater similarities (lower beta diversity) in the CWC fauna at upper bathyal depths and a distinct OMZ community compared to deeper depths. There was marked species turnover across the OMZ upper and lower boundaries, resulting in these three groupings. In addition, no coral species were observed to occur both above and below the OMZ (Fig. 4), suggesting that dissolved oxygen concentration is a critical control on coral species distribution on features in the eastern tropical Pacific Ocean, and may even by a barrier to gene flow between populations of CWC reinforcing allopatric speciation mechanisms (Rogers, 2000).

We did not find any differences in deep-water coral assemblages between oceanic water masses in this region but acknowledge that additional surveying effort in the shallowest (Equatorial Subsurface Water) and deeper water masses (North Pacific Deep Water and Circumpolar Deep Water) is warranted. In other basins, oceanographic water masses have been found to significantly influence community structure changes across bathyal depths (Ayress, De Deckker & Coles, 2004; Arantes et al., 2009; Quattrini et al., 2015; Radice et al., 2016; Auscavitch & Waller, 2017; Victorero et al., 2018; Auscavitch et al., 2020), leading to increased local (e.g. within-feature) and regional deep-water coral diversity. However, as discussed below, the water mass structure of the upper bathyal eastern tropical Pacific is complex and strongly influenced by equatorial ocean current structure in the area (Firing et al., 1998; Fiedler & Talley, 2006; Evans et al., 2020), which may make it difficult to assign characteristic assemblages to water masses at the present sampling resolution.

### The effects of OMZs on deep-water coral communities off Costa Rica

Deep-water coral assemblages were observed to change mostly as a function of dissolved oxygen changes in the water column and depth (Fig. 12). Coral abundances were slightly depressed across the upper boundary and within the OMZ and peaked along the lower boundary (Fig. 6), a pattern which has been observed in other taxa across both benthic and pelagic OMZ habitats (Gooday et al., 2010; Wishner et al., 2013). Likewise, species richness (S) and Shannon’s diversity index (H’) were observed to decline through the upper boundary of the OMZ and increased along the lower boundary (Fig. 7A, B), peaking at intermediate bathyal depths. As an indicator of beta diversity, the changes in species evenness across shallower depths (Fig. 7D) are consistent with high rates of species turnover across depths above and within the OMZ (Fig. 4). Higher evenness at mid- and lower bathyal depths is indicative of more equitable representation of coral species with increasing depth.

These observed patterns in faunal abundance and diversity peaks along the lower OMZ boundary are thought to be a result of three potential mechanisms: 1) the selective pressures that might exclude deep-water coral species from hypoxic or dysoxic conditions (Gooday et al., 2010; McClain & Hardy, 2010), 2) the potential for low oxygen tolerance provided by a metabolic offset associated with enhanced food supply along the lower OMZ (Rogers, 2000), and 3) the adaptation of sessile benthic organisms to natural variability in bottom oxygen concentration (Levin, 2003) changes along the lower OMZ boundary as a result of oceanographic processes (e.g. internal waves and tidal dynamics) (Hanz et al., 2019).

Several octocoral species were found to occur in the greatest abundances within 100 m of the lower boundary of the OMZ. For one sea pen, *Pennatula* cf. *inflata*, 96% of all records for this species occurred at Seamount 6 between a narrow depth range of 653-669 m (Supplementary Table 1). Additionally, 4 of the 5 large, branching paragorgiidae (since combined into the Coralliidae *sensu* McFadden et al., 2022) morphospecies identified here, consisting of both red and white color morphs, were narrowly distributed along the lower boundary of the OMZ (549-893 m). In total, 21% (778) of all coral occurrences were observed within +/- 100 m of the lower boundary (700 m) of the OMZ, in contrast to the 254 records +/-100 m of the upper OMZ boundary (300 m). In the eastern Pacific, the existence of a peak in suspended particulate organic carbon (POC) around the lower boundary of the OMZ near Volcano 7, off Mexico, has been identified as a potential driver of higher benthic megafauna abundance (Wishner et al., 1995).

Because organic matter degradation rates will be slower under hypoxia and anoxia, the OMZ may serve to deliver more and higher quality particulate organic matter to greater depths, and result in a fauna that is adapted to take advantage of this.

This study also gives us insights to coral taxa which are low-oxygen tolerant and are able to colonize OMZ depths. In general, stylasterids as a group, and *Stylaster* spp. specifically, were among the most broadly distributed taxa across the bathyal oxygen gradient and the most likely to be disproportionally abundant within the OMZ (Supplementary Table 4). This observation is particularly noteworthy since they can be an important habitat and structure-forming group in the deep sea (Häussermann & Försterra, 2007; Buhl Mortensen et al., 2010). Also within the OMZ, 304 coral observations across 10 species were made at oxygen concentrations under 5 µmol/L (371-582 m depth), primarily at Las Gemelas and Seamount 6. At the most extreme oxygen values <1 µmol/L, only one species was identified: Plexauridae sp. S0227-Q9, with 40 colonies between 495-512 m and only occurring at Seamount 6. However, it is important to note that ambient anoxic conditions measured on the seafloor are not static and may have some component of temporal variability relating to tidal dynamics, seasonal current variability, and internal waves (Hanz et al., 2019).

The observed distribution of coral assemblages across the Costa Rica seamounts likely reflects the compounding effects of multiple co-varying abiotic stressors rather than the influence of any single environmental variable. Above the OMZ (<300 m), adequate dissolved oxygen, warm temperatures, and supersaturated aragonite conditions together constitute a physiologically favorable environment for reef-building taxa including scleractinians and stylasterids, which require both oxygen for aerobic metabolism and skeletal deposition. Within the OMZ (300-700 m), corals are simultaneously exposed to severe hypoxia, approaching anoxia at the OMZ core, colder temperatures, and unfavorable conditions for skeletal aragonite precipitation. In this multi-stressor environment, only taxa with physiological or bio-mineralogical adaptations (e.g. Octocorallia and Antipatharia) would benefit owing to a resistance to both hypoxia and carbonate undersaturation. Below the OMZ (>700 m), oxygen partially recovers as NPDW advects from the north, and temperatures stabilize at near-bottom values around 4°C; however, Ω_arag_ remains persistently low, below 0.7, indicating that carbonate undersaturation is the dominant and unrelenting chemical stressor at bathyal depths.

The capacity of octocorals and antipatharians to dominate the bathyal zone despite persistent aragonite undersaturation likely reflects their proteinaceous or calcite-based skeletal chemistry (Cairns & MacIntyre, 1992), which is more favorable than aragonitic corals (Gomez et al., 2022). Together, these observations suggest that the ecological transition zones at the upper (∼300 m) and lower (∼700 m) OMZ boundaries represent multi-stressor ecotones rather than simple oxygen thresholds, and that future deoxygenation and ocean acidification may act synergistically to compress the depth range of calcifying taxa along the Eastern Tropical Pacific margin. Future work should aim to establish a more refined understanding of the distribution of species sensitive to aragonite saturation, temperature, and oxygen stressors and experimentally evaluate the species in a controlled laboratory setting.

### Insights to Pacific Deep-water Coral Biogeography

Cold-water coral species inventories in the eastern tropical Pacific are further improved by the results of this study and, consequently, these data provide an important context in which to evaluate proposed biogeographic provinces in the deep ocean. Bathyal depths across the area encompassing eastern Pacific latitudes from the Gulf of California to the continental margin off Peru, including the Cocos Ridge and Galapagos Islands, have previously been identified as a single biogeographic unit (BY7: Cocos Plate) (Watling et al., 2013). This province has been characterized as having a moderately high particulate organic carbon (POC) flux (4-6 g/m^-2^/y^-1^) to the seabed and relatively low dissolved oxygen concentration (1 mL L^-1^). It is our finding that the bathyal depths of Cocos Island and nearby seamounts host cosmopolitan, pan-Pacific, and possibly endemic species in this region.

Within the Primnoidae, *Calyptrophora clarki* Bayer, 1951 was observed between 896 -1138 m is known to be distributed across the Pacific basin at similar depths (Cairns, 2018b). This same species has been found off Hawaii (Cairns & Bayer, 2009) and was identified during explorations of the Phoenix Islands Protected Area and Northern Mariana Islands (Cairns, 2018b; Auscavitch et al., 2020). At the same time, two species of *Callogorgia* are also present in this area, one of which, *Callogorgia kinoshitai* Kükenthal, 1913, appears to be restricted to the deeper Costa Rica margin (specimens from cruises AT- 37; unpublished data) and was not encountered along the seamounts in this study, despite having been collected in the Galápagos archipelago (Cairns, 2018a). The other, *Callogorgia galapagensis* Cairns, 2018, initially described from the Galápagos is present throughout the study area (Las Gemelas, Seamount 6, 241-661 m) and has also been previously identified at Cocos Island between 628-656 m (Cairns, 2018a). In the genus *Narella*, one species encountered two times in this study and collected once (S0221-Q3), *Narella ambigua* Studer, 1894, has only ever been identified in this region of the tropical eastern Pacific in the Galapagos (Cairns, 2018a) and now at Seamount 5.5 between 1529-1548 m.

Additionally we extend the depth distribution and range of *Parastenella pomponiae* Cairns, 2018. Initially described from material collected in the Galápagos between 475-578 m (Cairns, 2018a), *Parastenella* cf. *pomponiae* was observed 19 times between 949-1398 m at Seamounts 5.5, 7, and 8. Additional collections from earlier Costa Rica margin cruises using the HOV *Alvin* (AT-37 and AT-42) also indicate its presence along the Costa Rica margin at Seamounts 1 (CR4925-1), 3 (CR4982-A1), and on the Quepos Plateau (CR4981-A2) down to 1752 m suggesting a truly wide depth distribution for this species in the region (Cairns et al., 2026).

Other octocoral taxa from the seamounts that provide additional biogeographic insight come from the Chrysogorgiidae and superfamily Pennatuloidea. Of the three *Chrysogorgia* species observed, two, *Chrysogorgia scintillans* Bayer and Stefani, 1988 and *Chrysogorgia midas* Cairns, 2018, were previously identified in the eastern Pacific from collections in the Galápagos (Cairns, 2018a). The most widespread, *Chrysogorgia scintillans* has a broad equatorial and North Pacific distribution (Cairns, 2018a). In contrast, *Chrysogorgia* cf. *midas* (this study), was identified at Seamounts 4, 5.5, and 7 between 866-1539 m, which represents a depth range extension since its description from material in the Galápagos (560-816 m) (Cairns, 2018a). Lacking from the study area were the chrysogorgiid genera *Metallogorgia* and *Iridogorgia*, often observed in seamount environments throughout Pacific Ocean (Watling et al., 2011; Pante & Watling, 2012; Pante et al., 2012; Auscavitch et al., 2020).

Pennatuloids, despite being represented by only 4 species, were locally abundant on some seamounts (Seamounts 4, 6, and 7) and were largely represented by two species, *Pennatula* cf. *inflata*, and *Anthoptilum* cf. *lithophilum*. The most abundant sea pen, *Anthoptilum* cf. *lithophilum*, noted for its ability to attach onto rocky hard substrate, appears to be widely distributed across the northeastern Pacific margin and Galápagos archipelago (Williams & Alderslade, 2011; Buglass et al., 2019). *Pennatula* cf. *inflata* has been reported globally (López-González, Gili & Williams, 2001; Schejter et al., 2018) as well as nearby in the Galápagos (Buglass et al., 2019). Both species were found to occur at similar depths but primarily in the lower OMZ and deeper (Fig. 4, Supplementary Table 1).

Stony corals were more well-represented by solitary species than colonial ones in this region. Of the 12 scleractinian species identified, 10 were solitary cup corals. The two colonial forms were *Madrepora oculata* (790-873 m), a cosmopolitan framework-forming species, and the shallower (180-320 m) dendrophylliid (Dendrophylliidae sp. S0224-Q1). Solitary species included *Desmophyllum dianthus* (574-783 m) and *Javania* sp. S0226-S8 (1217-1442 m) and several others of uncertain families (Supplementary Table 1). These early records indicate that stony coral fauna at bathyal depths in this region are likely to be composed of widespread species, but further examination of the many unidentified solitary individuals is required.

Finally, within the Antipatharia (black corals), one species observed in this area, *Lillipathes ritamariae* (913-1167 m), initially described from collections off Costa Rica (Opresko & Breedy, 2010), has since been observed to occur across the South Pacific and Southern Ocean with more recent records off Chile (Araya, et al., 2018), and as far west as New Zealand (Opresko, et al., 2014). Further examination of black coral collections from this and other recent expeditions along the Costa Rica margin are needed to obtain better species-level identifications in support of biogeographic analyses.

The complex water mass and equatorial current structure of the equatorial eastern Pacific provides a potential mechanism for the observation of widely distributed Pacific species in the region of the Costa Rica Dome and along the seamounts. At upper mid-bathyal depths (200-1500 m), Equatorial Pacific Intermediate Water (EqPIW), the largest water mass explored in this study, is sourced from Antarctic Intermediate Water (AAIW) extending northward from the South Pacific and North Pacific Intermediate Water (NPIW) and North Pacific Deep Water (NPDW) from the north (Bostock et al., 2010). The mixing of these geographically extensive water masses into EqPIW provides a potential corridor for distribution of bathyal coral species from the northern and southern hemispheres in the eastern equatorial Pacific. Furthermore, observed peaks in coral diversity at intermediate depths (1200-1500 m) (Fig. 7), suggest that elevated diversity at intermediate depths may be a result of the mixing of multiple deep water masses and overlapping depth ranges of upper and mid-bathyal taxa. Below 900 m, the presence of widely distributed species with increasing depth appears to be characteristic of deeper ocean water masses, in this case NPDW.

On larger scales, the role that shallower currents have in species distributions across the Pacific Ocean, and particularly laterally along the equator, should not be overlooked as a means of biogeographic connectivity at upper bathyal depths. The flow of the Equatorial Undercurrent (EUC) provides rapid transport eastward along the equator eastward from the Western and Central Pacific directly to the equatorial eastern Pacific at depths between 150-300 m (Taft et al., 1974; Tsuchiya, 1981; Evans et al., 2020). In addition to being a potential vector for conveying rapid (0.9-1 m s^-1^) transport of planktonic coral larvae, this current supplies oxygen to upper bathyal depths are along the equator due to transport of more well-oxygenated water from the western Pacific along the EUC (Tsuchiya et al., 1989; Fiedler & Talley, 2006; Stramma et al., 2010a). Below this (∼500 m), a return current, the Equatorial Intermediate Current (EIC), carries a comparatively oxygen-poor water westward (Stramma et al., 2010a).

As an important transport mechanism for dissolved oxygen, as well as seamount coral larvae, these currents represent potentially important dispersal vectors along vast expanses of the equatorial Pacific. The volume of oxygen that the EUC delivers to the equatorial eastern Pacific bathyal depths may even offset the effects of projected OMZ expansion in the region (Stramma et al., 2010a). As additional deep-water species distribution data continues to emerge from across the Pacific (Friedlander et al., 2019; Giddens et al., 2019; Kennedy et al., 2019; Auscavitch et al., 2020; López-Garrido et al., 2020), future biogeographic studies should continue to take into account physical oceanographic features, like water masses and major ocean currents, for identifying potential dispersal and connectivity pathways across the vast expanse of the Pacific Ocean Basin.

### Cold-water Corals, Ocean Deoxygenation, and Synergistic Multistressor Effects

This work provides baseline metrics of deep-water coral biodiversity on several previously unexplored Costa Rican seamounts, which are vital to understanding the effects that projected expansion of OMZs and ocean deoxygenation (Stramma et al., 2008; Gilly et al., 2013) will have on benthic communities. Measurements for the eastern tropical Pacific indicate expansion and intensification over the past half century with oxygen loss between 300-700 m as 0.09-0.34 µmol kg^-1^ y^-1^ (Stramma et al., 2008). With tropical oxygen minimum zones expanding horizontally and vertically (Stramma et al., 2010b), the strongest effects of ocean deoxygenation are likely to occur at upper bathyal and bathypelagic environments (Stramma et al., 2010b; Wishner et al., 2018; Wishner, Seibel & Outram, 2020). Effects on CWC distributions may result in compression of upper bathyal species distributions, particularly those with narrow depth ranges along the upper boundary of the OMZ and shallower (Fig. 4). However, those species with broader depth distributions, such as those inhabiting depths below the OMZ, may be more resilient to such expansions (McClain & Hardy, 2010).

The physiological tolerances of framework-forming stony coral species specifically may limit their distribution to more well-oxygenated waters (Dodds et al., 2007; Lunden et al., 2014), but in some cases they have been observed thriving within hypoxic conditions (Hebbeln et al., 2020). Across all sites we noted a poor representation of framework-forming scleractinian species. One cosmopolitan deep-water species, *Madrepora oculata*, was observed 8 times between 790-873 m and only at Seamount 5.5. A second colonial species, Dendrophylliidae sp. S0224-Q1, was also found to be restricted largely above the depth of the OMZ (180-320 m) and only at the sites furthest from the continental margin (Las Gemelas, Cocos South). As with many other coral taxa, colonial scleractinian species are important for their function as ecosystem engineers, producing hard structures that can lead to increased biodiversity in the deep ocean (Roberts et al., 2009; Buhl Mortensen et al., 2010). As a result of deoxygenation, the displacement or loss of coral fauna and the structures they produce, combined with other climate change-driven stressors (e.g. deep-ocean warming (Brito-Morales et al., 2020), changes in surface productivity (Ovsepyan, Ivanova & Murdmaa, 2018), and ocean acidification (Perez et al., 2018)), will likely have manifold effects on the diversity of life supported within the deep benthic communities.

## Acknowledgements

We would like to extend our thanks to the Schmidt Ocean Institute, as well as the science party and crew of the R/V *Falkor* on FK190106: *Costa Rican Deep-sea Connections*. The ability to complete this manuscript was funded, in part, by a Temple University Dissertation Completion Grant to S. Auscavitch. Thanks also to Stephen Cairns (National Museum of Natural History, Smithsonian Institution) for taxonomic assistance in identifying voucher material. Also, special thanks to Salome Buglass for sharing additional reference material on deep-sea corals from the Galápagos Islands.

