## Supplementary Table 2 for "Oxygen Minimum Zones Drive Changes in Deep-sea Coral Species Distribution, Diversity, and Community Structure on Seamounts in the Eastern Tropical Pacific"

Supplementary Table 2: One-way pairwise ANOSIM comparisons between coral assemblages above, within, and below the OMZ along the Costa Rica seamounts. Values indicated are R-statistics with p-values indicated in parentheses. Values in bold were observed to be significant at or below the p=0.05 level.

| Factor | Above OMZ | Within OMZ | Below OMZ |
| --- | --- | --- | --- |
| Above OMZ |  |  |  |
| Within OMZ | 0.144  (0.132) |  |  |
| Below OMZ | **0.683**  **(0.002)** | **0.587**  **(0.001)** |  |
