## Supplementary Table 3 for "Oxygen Minimum Zones Drive Changes in Deep-sea Coral Species Distribution, Diversity, and Community Structure on Seamounts in the Eastern Tropical Pacific"

Supplementary Table S3: One-way ANOSIM pairwise comparisons among water mass factors. Values indicated are R-statistics with p-values indicated in parentheses. Water masses are indicated by the following abbreviations: Equatorial Subsurface Water (EqSSW), Equatorial Pacific Intermediate Water (EqPIW), North Pacific Deep Water (NPDW).

| Water Mass | EqSSW | EqPIW | NPDW |
| --- | --- | --- | --- |
| EqSSW |  |  |  |
| EqPIW | 0.27  (0.053) |  |  |
| NPDW |  | 0.028  (0.50) |  |
