## Supplementary Table 4 for "Oxygen Minimum Zones Drive Changes in Deep-sea Coral Species Distribution, Diversity, and Community Structure on Seamounts in the Eastern Tropical Pacific"

Supplementary Table 3: Similarity percentage analysis results.

Parameters

Resemblance: S17 Bray-Curtis similarity

Cut off for low contributions: 70.00%

Group Below OMZ

Average similarity: 19.76

Species	Av.Abund	Av.Sim	Sim/SD	Contrib%	Cum.%

Hemicorallium cf. imperiale	1.24	4.53	0.67	22.92	22.92

Isididae sp. S0226-Q3	0.87	3.20	0.36	16.22	39.13

Bathypathes sp. S0226-Q9	0.85	2.75	0.47	13.94	53.07

Crypthelia sp. S0221-S8	0.90	2.54	0.42	12.86	65.93

Aquaumbra klapferi	0.56	1.11	0.22	5.59	71.52

Group OMZ

Average similarity: 12.00

Species	Av.Abund	Av.Sim	Sim/SD	Contrib%	Cum.%

Stylaster sp. S0224-Q3	1.10	3.93	0.42	32.75	32.75

Paragorgiidae sp. S0223-Q4	0.63	2.09	0.44	17.42	50.17

Scleractinia sp. 4	0.64	1.43	0.32	11.96	62.12

Plexauridae sp. S0227-Q9	0.55	0.95	0.13	7.88	70.01

Group Above OMZ

Average similarity: 31.71

Species	Av.Abund	Av.Sim	Sim/SD	Contrib%	Cum.%

Stylaster sp. 2	2.02	12.69	0.58	40.02	40.02

Dendrophylliidae sp. S0224-Q1	1.53	11.91	1.94	37.57	77.59

Groups Below OMZ & OMZ

Average dissimilarity = 98.57

Groups Below OMZ & Above OMZ

Average dissimilarity = 100.00

Groups OMZ & Above OMZ

Average dissimilarity = 91.41
