## Supplementary Table 5 for "Oxygen Minimum Zones Drive Changes in Deep-sea Coral Species Distribution, Diversity, and Community Structure on Seamounts in the Eastern Tropical Pacific"

Supplementary Table 5: Output data of the DistLM and dbRDA analysis.

DistLM

Distance based linear models

Resemblance worksheet

Data type: Similarity

Selection: All

Standardise Samples by Total

Transform: Fourth root

Resemblance: S17 Bray-Curtis similarity

Predictor variables worksheet

Name: Data15

Data type: Environmental

Sample selection: All

Variable selection: All

Normalise

Selection criterion: AIC

Selection procedure: Specified

VARIABLES

1	Depth	Trial

2	Temperature	Trial

3	Salinity	Trial

4	O2	Trial

5	Ωarag	Trial

Total SS(trace): 1.5925E5

MARGINAL TESTS

Variable	SS(trace)	Pseudo-F	P	Prop.

Depth	25693	7.1178	0.001	0.16134

Temp	23512	6.4091	0.001	0.14764

Salinity	17066	4.4411	0.001	0.10717

O2	22811	6.1858	0.001	0.14324

Ωarag	16388	4.2444	0.001	0.10291

res.df: 37

SEQUENTIAL TESTS

Variable	AIC	SS(trace)	Pseudo-F	P	Prop. Cumul. res.df

+Depth	321.41	25693	7.1178	0.001	0.16134	0.16134	37

+Temperature 319.45	12881	3.8428	0.001	8.0888E-2	0.24222	36

+Salinity	318.6	8527.8	2.6614	0.001	5.355E-2	0.29577	35

+O2	318.39	6160.8	1.9763	0.012	3.8686E-2	0.33446	34

+Ωarag	319.13	3378.1	1.0864	0.363	2.1213E-2	0.35567	33

Specified solution

AIC	R^2	RSS	No.Vars Selections

319.13	0.35567 1.0261E5	5 All

Percentage of variation explained by individual axes

% explained variation	% explained variation

out of fitted model	out of total variation

Axis	Individual Cumulative	Individual Cumulative

1	50.31	50.31	17.89	17.89

2	25.79	76.1	9.17	27.07

3	10.05	86.15	3.58	30.64

4	9.2	95.35	3.27	33.91

5	4.65	100	1.65	35.57

dbRDA coordinate scores

Sample	dbRDA1	dbRDA2	dbRDA3	dbRDA4	dbRDA5

S0221.700-800	10.994	17.615	-4.7546	-6.3608E-2	-2.7666

S0221.800-900	0.16956	14.503	-10.768	6.9602	-2.7142

S0221.900-1000	-17.307	4.4399	-9.7296	8.1237	3.2562

S0221.1000-1100	-24.078	-3.631	-8.8643	3.4819	11.44

S0221.1100-1200	-24.077	-0.16174	-5.026	-0.81993	-3.0533

S0221.1200-1300	-31.485	-6.434	-2.3348	-4.7222	-0.77104

S0221.1300-1400	-35.631	-5.5576	-0.35629	-5.1974	-12.243

S0221.1400-1500	-33.16	-9.8404	13.793	-19.425	-5.7345

S0221.1500-1600	-42.732	-18.866	11.805	-21.274	1.1367

S0222.800-900	-17.616	8.5938	-14.672	24.047	6.3188

S0222.900-1000	-14.159	9.0861	-6.1014	16.522	6.8053

S0222.1000-1100	-23.829	-2.0189	-6.456	5.7095	11.988

S0222.1100-1200	-28.418	-0.59945	-4.4199	6.5255	0.79287

S0222.1200-1300	-35.449	-7.0948	-4.3343	3.1997	3.2601

S0223.500-600	34.245	12.42	1.1658	-8.0278	2.7776

S0223.600-700	35.614	19.445	9.6532	-10.416	1.6485

S0223.700-800	20.835	23.458	13.22	1.6853	2.2145

S0223.800-900	14.049	25.508	16.775	7.0134	-0.34402

S0223.900-1000	4.6272	22.112	18.347	10.082	3.6245

S0224.100-200	31.683	-44.204	33.948	39.266	-2.5109

S0224.200-300	30.119	-43.189	-1.9825	8.1865	-12.787

S0224.300-400	33.566	-44.59	-1.648	-12.906	25.763

S0224.500-600	26.724	17.985	-4.9432	1.8702	-2.6622

S0225.200-300	25.431	-47.202	-14.991	3.4122	-17.945

S0225.300-400	31.09	-35.033	-16.174	-15.422	6.315

S0225.400-500	44.546	-0.50621	3.9923	-18.972	1.0538

S0225.500-600	37.251	11.579	7.7472	-7.9546	7.5249

S0225.600-700	30.33	19.402	8.8871	-0.88618	5.7374

S0226.1200-1300	-38.938	-7.9982	3.9524	5.9106	6.8397

S0226.1300-1400	-32.662	8.0609E-2	11.604	2.9592	-7.8067

S0226.1400-1500	-22.828	0.86829	30.041	-13.048	-3.4612

S0227.400-500	32.03	17.663	-13.932	0.11723	-19.306

S0227.500-600	23.838	10.058	-15.618	-1.8455	-0.4232

S0227.600-700	12.326	12.748	-18.193	3.0837	-1.207

S0228.800-900	0.96268	14.808	-3.8459	2.4609	-4.0875

S0228.900-1000	-4.6754	10.6	-3.4449	1.8324	-9.4957E-2

S0228.1000-1100	-19.935	-1.5912	-11.215	0.4089	8.2654

S0228.1100-1200	-14.925	4.667	-1.8087	-8.6127	-8.807

S0228.1200-1300	-18.525	0.87807	0.68136	-13.265	-8.0383

Relationships between dbRDA coordinate axes and orthonormal X variables

(multiple partial correlations)

Variable	dbRDA1	dbRDA2	dbRDA3	dbRDA4	dbRDA5

Depth	-0.458	-0.036	0.320	-0.755	-0.342

Temperature	0.658	-0.130	0.710	-0.159	0.146

Salinity	0.011	-0.899	-0.276	-0.204	0.272

O2	-0.571	-0.243	0.563	0.516	0.177

Ωarag	0.178	-0.338	0.022	0.312	-0.870

Weights

(Coefficients for linear combinations of X's in the formation of dbRDA coordinates)

Variable	dbRDA1	dbRDA2	dbRDA3	dbRDA4	dbRDA5

Depth	72.598	39.761	99.424	-70.077	-32.427

Temperature	179.42	107.99	228.43	-66.018	-14.908

Salinity	-110.04	-110.91	-138.75	9.3877	41.186

O2	7.1526	14.388	37.579	22.542	14.442

Ωarag	12.398	26.366	6.1065	7.3342	-49.949
